# Biphasic inflammation control by dedifferentiated fibroblasts enables axon regeneration after spinal cord injury in zebrafish

**DOI:** 10.1101/2025.01.27.635043

**Authors:** Nora John, Thomas Fleming, Julia Kolb, Olga Lyraki, Sebastián Vásquez-Sepúlveda, Asha Parmar, Kyoohyun Kim, Maria Tarczewska, Pooja Gupta, Kanwarpal Singh, Federico Marini, Sumeet Pal Singh, Sven Falk, Kristian Franze, Jochen Guck, Daniel Wehner

**Affiliations:** Max Planck Institute for the Science of Light, Erlangen, Germany; Max-Planck-Zentrum für Physik und Medizin, Erlangen, Germany; Department of Biology, Animal Physiology, Friedrich-Alexander-Universität Erlangen-Nürnberg, Erlangen, Germany; Institute of Medical Physics and Microtissue Engineering, Friedrich-Alexander-Universität Erlangen-Nürnberg, Erlangen, Germany; Department of Physics, Friedrich-Alexander-Universität Erlangen-Nürnberg, Erlangen, Germany; Department of Stem Cell Biology, University Hospital Erlangen, Friedrich-Alexander-Universität Erlangen-Nürnberg, Erlangen, Germany; Department of Electrical and Computer Engineering, McMaster University, Hamilton, Canada; Research Center for Immunotherapy (FZI), Mainz, Germany; Institute of Medical Biostatistics, Epidemiology and Informatics, University Medical Center of the Johannes Gutenberg-University, Mainz, Germany; IRIBHM, Université libre de Bruxelles, Brussels, Belgium; Institute of Biochemistry, Friedrich-Alexander-Universität Erlangen-Nürnberg, Erlangen, Germany; Department of Physiology, Development and Neuroscience, University of Cambridge, Cambridge, United Kingdom

**Keywords:** Zebrafish, spinal cord injury, axon regeneration, inflammation, fibroblasts, dedifferentiation, neutrophils, macrophages, single-cell transcriptomics, scar, Brillouin microscopy, optical coherence tomography, tissue mechanics, AFM-based nanoindentation

## Abstract

Fibrosis and persistent inflammation are interconnected processes that inhibit axon regeneration in the mammalian central nervous system (CNS). In zebrafish, by contrast, fibroblast-derived extracellular matrix deposition and inflammation facilitate regeneration. However, the regulatory cross-talk between fibroblasts and the innate immune system in the regenerating CNS is not understood. Here, we show that zebrafish fibroblasts possess a dual role in inducing and subsequently resolving inflammation, which are both essential for regeneration. We identify a transient, injury-specific *cthrc1a*^+^ fibroblast state with an inflammation-associated, less differentiated, and non-fibrotic profile. Induction of this fibroblast state precedes and contributes to the initiation of the inflammatory response. At the peak of neutrophil influx, *cthrc1a*^+^ fibroblasts coordinate the resolution of inflammation. Disruption of these inflammation dynamics inhibits axon regeneration and alters the mechano-structural properties of the lesion environment. This establishes the biphasic inflammation control by dedifferentiated fibroblasts as a pivotal mechanism for CNS regeneration.

**ONE SENTENCE SUMMARY:** Dedifferentiated fibroblasts sequentially induce and resolve neutrophil-driven inflammation through cytokine release to facilitate axon regeneration after spinal cord injury in zebrafish.

**HIGHLIGHTS:**

- Time-resolved single-cell transcriptomics of zebrafish spinal cord regeneration.
- Spinal cord injury induces fibroblast dedifferentiation.
- Dedifferentiated fibroblasts sequentially induce and resolve inflammation.
- Dysregulation of inflammation dynamics alters mechano-structural tissue properties.

## INTRODUCTION

Spinal cord injury (SCI) results in sensorimotor and autonomic dysfunction. In humans and other adult mammals, these functional deficits persist because of the failure of the severed axons to regenerate across the lesion ^1^. Prolonged inflammation and the formation of fibrotic scar tissue—two intertwined processes—are key contributors to regeneration failure ^2–6^. Fibrous scarring in the central nervous system (CNS) refers to the excessive deposition of extracellular matrix (ECM), mainly by fibroblasts, that exhibits biochemical and mechanical (viscoelastic) properties adverse to axon growth ^7–11^. This process has a reciprocal relationship with inflammation, as injury-induced immune cell infiltration promotes scarring, which in turn intensifies and prolongs the immune response, impeding resolution of inflammation ^2,5,12^.

In zebrafish, unlike mammals, the immune response and fibroblast-derived injury ECM promote axon regeneration after SCI ^11,13–19^. Emerging evidence suggests a requirement for tight regulatory control over these processes to facilitate the differential regenerative outcomes between mammals and zebrafish. For example, neutrophil invasion is rapidly resolved, curtailing the acute inflammatory phase ^15,17,20^. Additionally, infiltrating fibroblasts selectively cease production of inhibitory ECM proteins, including members of the small leucine-rich proteoglycan family, which deteriorate the mechanical properties of the lesion environment ^11,13^. However, little is known about the regulatory interactions between fibroblasts and innate immune cells in the injured zebrafish spinal cord, as these cell populations have primarily been studied separately. Furthermore, while fibroblast heterogeneity is increasingly recognized in the injured mammalian CNS, the diversity during zebrafish spinal cord regeneration has yet to be explored ^21–24^.

Here, we employed time-resolved single-cell transcriptomics to identify fibroblast populations and their interactions with effector cells of the innate immunity after SCI. We identify a transient population of *cthrc1a*^+^ fibroblasts is characterized by a less differentiated, non-fibrotic profile, which possesses dual functions in modulating inflammation. *cthrc1a*^+^ fibroblasts release both pro-inflammatory and immunosuppressive cytokines to induce and subsequently resolve the acute inflammatory phase, establishing a lesion environment with regeneration-permissive properties. Our findings demonstrate that fibroblasts biphasically control inflammation to enable axon regeneration in the zebrafish CNS.

## RESULTS

### Time-resolved single-cell transcriptomics identify reactive fibroblast states

To explore fibroblast–immune cell interactions directing axon regeneration, we performed single-cell RNA sequencing (scRNA-seq) following spinal cord transection in larval zebrafish at 3–4 days post-fertilization (dpf). In this model, which shares key cellular responses with adult zebrafish after SCI, axon regeneration and recovery of swimming capacity are achieved within 48 hours post-lesion (hpl) (Fig. 1a–c) ^11,13–15,25–33^. To capture the entire regeneration time course with high temporal resolution, we dissociated trunk tissues containing the lesion site at 6, 12, 24, and 48 hpl and unlesioned controls (Fig. 1c). After quality control, we obtained 91,566 cells across two biological replicates for each condition for downstream analysis. Unsupervised leiden clustering of the filtered cells resulted in 27 major clusters, which were annotated based on known marker genes and encompassed the expected cell types, including all neural cells of the spinal cord (Fig. 1d; Supplementary Fig. 1; Supplementary File 1). These data can be explored at https://shiny.mpl.mpg.de/wehner_lab/spinal_cord_regeneration_atlas/.

**Figure 1.**
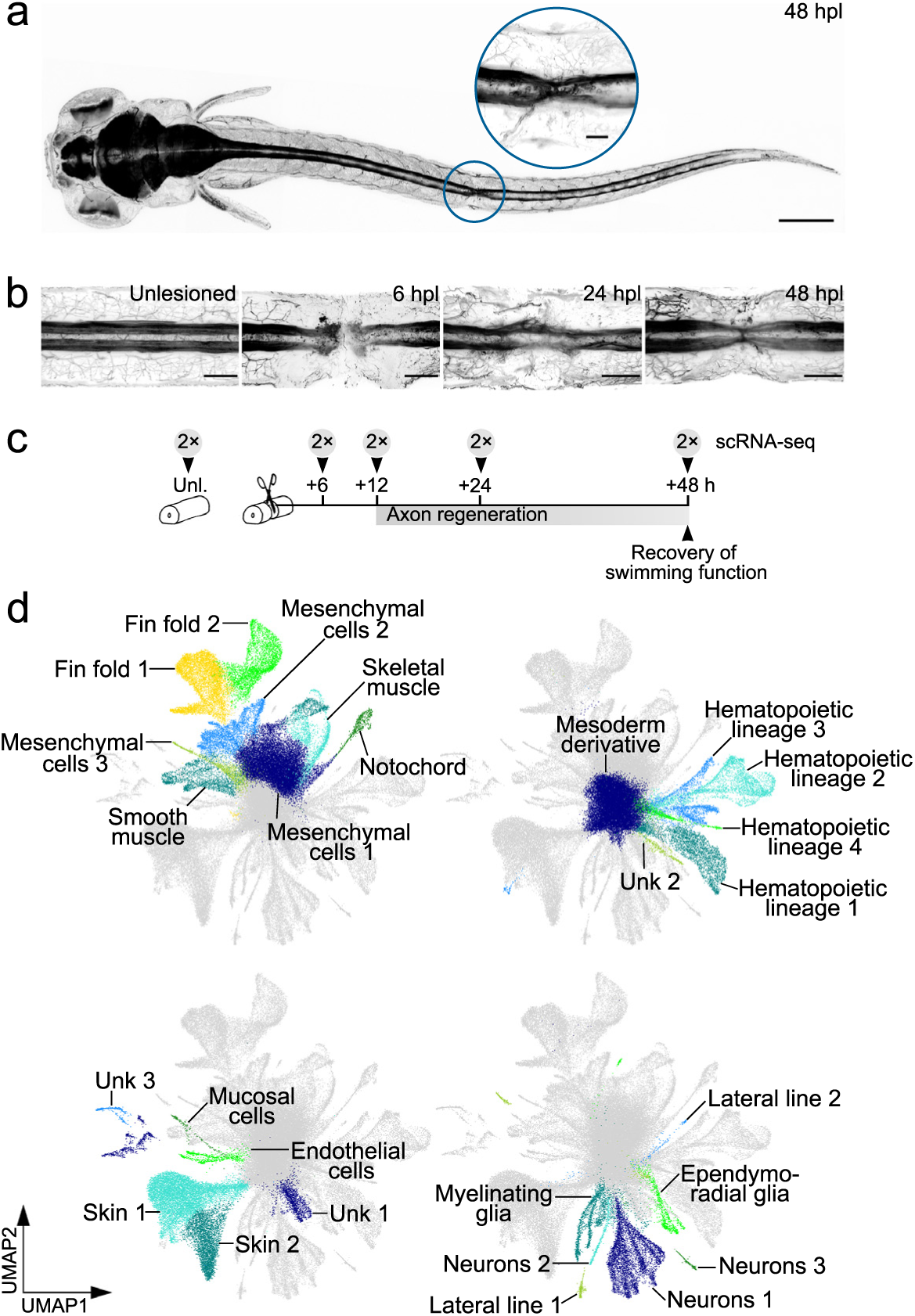
Single-cell transcriptomic analysis of spinal cord regeneration in zebrafish. **a)** *elavl3*:GFP-F transgenic zebrafish larva at 48 hours after spinal cord transection (hpl). Neurons and neurites are indicated in black. Circle indicates the dimension of trunk tissue analyzed by scRNA-seq. 3D reconstruction; dorsal view (rostral is left). **b)** Time course of axon regeneration after SCI in *elavl3*:GFP-F transgenic zebrafish larvae. Neurons and white matter tracts are indicated in black. 3D reconstruction; dorsal view (rostral is left). **c)** Timeline of axon regeneration and recovery of swimming function after SCI in larval zebrafish. Time points of tissue collection for scRNA-seq analysis and number of biological replicates is indicated. **d)** UMAP representation of the post filtering scRNA-seq data and clustering results. **a**–**d)** Abbreviations: h, hours; hpl, hours post-lesion; UMAP, uniform manifold approximation and projection; SCI, spinal cord injury; unk, unknown; unl, unlesioned. Scale bars: 250 µm (**a**), 50 µm (**b**), 25 µm (**a**, inset).

To identify reactive fibroblast populations, we first filtered for clusters with a 3-fold higher expression score of core matrisome genes (encoding collagens, proteoglycans, and glycoproteins) compared to the mean score of all cells (Supplementary File 2) ^34^. This analysis identified seven fibroblast-like clusters: mesenchymal cells 1–3, fin fold 1 and 2, smooth muscle, and notochord (Fig. 2a; Supplementary Fig. 2a). Second-level clustering of all fibroblast-like cells revealed 25 transcriptionally distinct subclusters, seven of which appeared transiently at the 6/12/24 hpl time points and contained few or no cells from unlesioned and 48 hpl samples, when axon regeneration has completed (Fig. 2b; Supplementary Fig. 2b) ^15^. These injury-specific transcriptional states were present in five of the seven fibroblast-like clusters and characterized by the expression of *cxcl18a.1* (fin fold 1), *cd44* (fin fold 1), *col8a1a* (notochord), *cthrc1a* (mesenchymal cells 2), *desmb* (smooth muscle), *foxl1* (smooth muscle), or *fbln1* (mesenchymal cells 3) (Fig. 2b). The injury-specific fibroblast populations demonstrated distinct transcriptional profiles with variations in the expression levels of matrisome genes, including those previously identified as promoters of axon regeneration in the zebrafish and mammalian CNS: *hbegfa* ^31^, *ntf3* ^35^, *col12a1a/b* ^14^*, crlf1a* ^36^*, cthrc1a* ^13^, *thbs1a/b* ^37,38^*, ctgfa* ^28^*, fn1a/b* ^39^*, tnc* ^40,41^, *tgfb3* ^20^, *sparc* ^20^, *dcn* ^42^, and *mmp2* ^43^ (Supplementary File 3; Fig. 2c,d). Notably, *cthrc1a*^+^ fibroblasts expressed the highest number of these genes, suggesting a pro-regenerative role for this cell population (Fig. 2d). *In situ* hybridization (ISH) revealed that *cthrc1a* expression was upregulated in regions surrounding the spinal cord and at myotendinous junctions, both near and distant from the lesion core, at 6 hpl—known fibroblast niches (Fig. 2e; Supplementary Fig. 2c) ^13,14,44^. By 24 hpl, *cthrc1a* transcripts localized to the lesion core (Fig. 2e). Based on these observations, we focused our subsequent analyses on *cthrc1a*^+^ fibroblasts.

**Figure 2.**
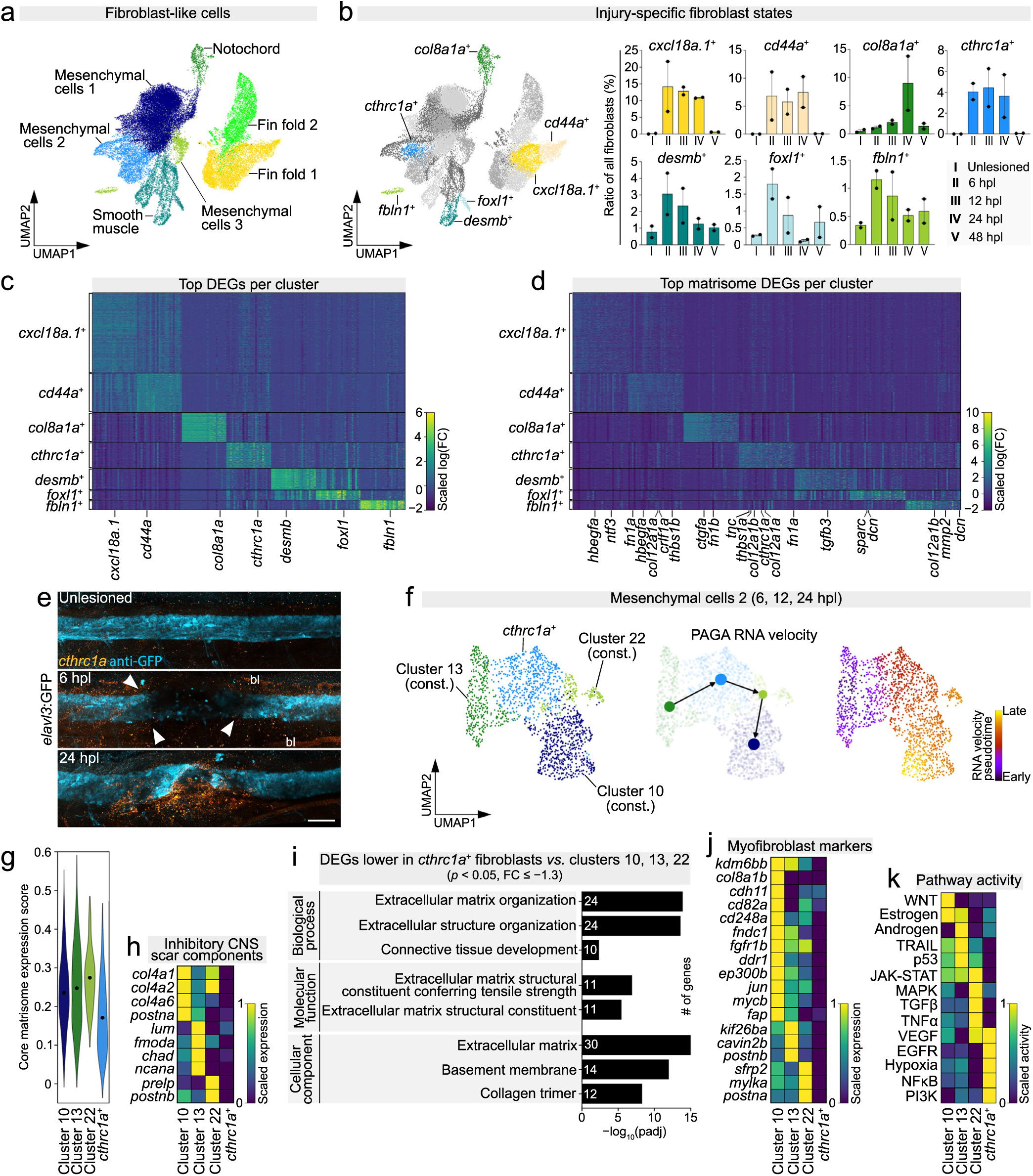
Dedifferentiation leads to the transient formation of *cthrc1a*^+^ fibroblasts after SCI. **a)** UMAP representation of all fibroblast-like cell clusters identified in Supplementary Fig. 2a. **b)** UMAP representation and cell number dynamics over all time points of injury-specific fibroblast subclusters. Each data point represents one independent biological replicate, normalized to the total number of fibroblasts per sample. Data are means ± SEM. **c and d)** Injury-specific fibroblast populations are transcriptionally heterogenous. Scaled expression heatmap of the top 25 DEGs (**c**) and top 31 matrisome DEGs (**d**) in each injury-specific fibroblast subcluster. Matrisome DEGs with known pro-regenerative roles are indicated in (**d**). **e)** mRNA expression of *cthrc1a* (orange) is upregulated in myotendinous junctions at the wound margin at 6 hpl (arrowheads) and localizes to the lesion core at 24 hpl. Images shown are maximum intensity projections of the unlesioned spinal cord (cyan) or lesion site (lateral view; rostral is left). *n* ≥ 10 for each time point. Scale bars: 50 µm. **f)** RNA velocity analysis of the mesenchymal cells 2 subclusters from 6, 12, and 24 hpl time points. **g)** Expression score of core matrisome genes of mesenchymal cells 2 subclusters. Data points represent means. **h)** Scaled expression of genes encoding inhibitory components of the mammalian CNS scar. **i)** Gene Ontology analysis of DEGs (FC ≤ −1.3, *p* < 0.05) between *cthrc1a*^+^ fibroblasts and constant fibroblasts. Fisher’s Exact test. **j)** Scaled expression of myofibroblast markers and genes associated with myofibroblast differentiation or function in mesenchymal cells 2 subclusters. **k)** Inferred activity of indicated signaling pathways in mesenchymal cells 2 subclusters. **a**–**k)** Abbreviations: bl, blood; const., constant; DEG, differentially expressed gene; FC, fold change; hpl, hours post-lesion; padj, *p*-value adjusted; PAGA, partition-based graph abstraction; UMAP, uniform manifold approximation and projection.

### Injury-specific cthrc1a^+^ fibroblasts are less differentiated

To predict the cellular dynamics of *cthrc1a*^+^ fibroblasts, we computed RNA velocity for the four subclusters of mesenchymal cells 2 (*cthrc1a*^+^, #10, #13, and #22) from the 6, 12, and 24 hpl time points. Clusters #10, #13, and #22 exhibited a relatively constant number of cells across all time points and will henceforth be referred to as constant fibroblasts (Supplementary Fig. 2b). Cell trajectories were visualized using uniform manifold approximation and projection (UMAPs) with pseudotime color code or partition-based graph abstraction (PAGA) ^45,46^. We found a unidirectional flow originating from cluster #13 toward *cthrc1a*^+^ fibroblasts, sequentially followed by clusters #22 and #10, indicating that *cthrc1a*^+^ fibroblasts derive from and subsequently transition to constant fibroblasts (Fig. 2f). Cell cycle gene expression analysis suggested an enhanced proliferation of *cthrc1a*^+^ fibroblasts at the 12 and 24 hpl time points (Supplementary Fig. 2d; Supplementary File 4). Interestingly, *cthrc1a*^+^ fibroblasts exhibited a 27–38% lower mean expression score for core matrisome genes than individual constant fibroblast clusters, suggesting a less differentiated cellular state (Fig. 2g). The core matrisome genes with reduced expression in *cthrc1a*^+^ fibroblasts included those associated with the inhibitory properties of the mammalian CNS scar, such as small leucine-rich proteoglycans (*lum*, *fmoda*, *chad*, *prelp*) ^11^, chondroitin sulfate proteoglycans (*ncana*) ^47^, periostin (*postna/b*) ^48^, and the basal lamina component collagen type IV (*col4a1*, *col4a2*, *col4a6*) ^49,50^ (Fig. 2h). Gene Ontology analysis of differentially expressed genes (DEGs) further supported reduced ECM secretion by *cthrc1a*^+^ fibroblasts compared to constant fibroblasts (Fig. 2i). Moreover, several myofibroblast markers and genes associated with myofibroblast differentiation/function exhibited lower expression in *cthrc1a*^+^ fibroblasts compared to constant fibroblasts (Fig. 2j) ^51–69^. Pathway activity inference analysis predicted reduced activity of Wnt and TGFβ signaling—pathways known to stimulate (myo)fibroblast differentiation—in *cthrc1a*^+^ fibroblasts compared to individual constant fibroblast clusters (Fig. 2k) ^70,71^. Altogether, these data indicate that *cthrc1a*^+^ fibroblasts represent a less differentiated, non-fibrotic cell population, which transiently arises through dedifferentiation in response to SCI.

### cthrc1a^+^ fibroblasts have an inflammation-associated profile

We noticed that several signaling pathways associated with inflammatory processes were predicted to exhibit high activity in *cthrc1a*^+^ fibroblasts (e.g., NFκB) and other mesenchymal cells 2 subclusters (e.g., TNFα) (Fig. 2j). Analyzing the expression of genes coding for cytokines and other inflammation-related secreted factors revealed that mesenchymal cells 2 had the second highest score across all major clusters (Fig. 3a; Supplementary File 5). Among the mesenchymal cells 2 subclusters, *cthrc1a*^+^ fibroblasts exhibited the highest expression levels of several cytokines with pro-inflammatory (e.g., *cxcl8b.1*, *cxcl11.5*, *cxcl18b*, *csf1a*, *il34*) or anti-inflammatory (e.g., *il11a*, *il11*, *tgfb1a*, *tgfb1b*, *tgfb2*) roles (Fig. 3a). To investigate the relevance of the inflammation-related profile of *cthrc1a*^+^ fibroblasts for axon regeneration, we focused on the NFκB pathway—a key mediator of inflammatory responses through cytokine expression regulation (Fig. 2k) ^72^. To monitor NFκB pathway activity in mesenchymal cells 2, we used *8xHs.NFκB*:GFP reporter zebrafish, combined with *pdgfrb*:memKillerRed transgenics to label *pdgfrb*^+^ cells. High *pdgfrb* and *cthrc1a* expression is largely restricted to mesenchymal cells 2, with *pdgfrb* expression virtually absent in hematopoietic lineage cells, making the *pdgfrb* promoter well suited for monitoring mesenchymal cells 2 that give rise to *cthrc1a*^+^ fibroblasts (Fig. 3b,c; Supplementary Fig. 3a). In unlesioned animals, strong NFκB reporter activity was detected in the caudal hematopoietic tissue, likely representing neutrophils (Supplementary Fig. 3b). At 6–12 hpl, NFκB reporter activity localized to the lesion core in memKillerRed^−^ putative neutrophils and was additionally detectable in memKillerRed^+^ mesenchymal cells 2 at the wound margin, consistent with the scRNA-seq analysis (Fig. 3d). To probe the role of the NFκB pathway in *cthrc1a*^+^ fibroblasts for axon regeneration, we inhibited NFκB transcriptional activity using JSH-23. Axon regeneration (measured by axonal bridge thickness ^15^) was reduced by 44% in JSH-23 treated animals at 48 hpl compared to DMSO controls (Supplementary Fig. 3c). Next, we used *pdgfrb*:CreERT2;*hs*:^loxP^Luc^loxP^IκBSR transgenic animals to induce expression of an NFκB signaling inhibitor (IκB super repressor [IκBSR]) in mesenchymal cells 2 after tamoxifen-induced Cre-mediated recombination (Supplementary Fig. 3d). This led to a 43% reduction in axonal bridge thickness compared to controls at 48 hpl (Fig. 3e). Thus, NFκB signaling in mesenchymal cells 2 is required for axon regeneration. Together, these data indicate that *cthrc1a*^+^ fibroblasts have an inflammation-associated profile that promotes CNS regeneration.

**Figure 3.**
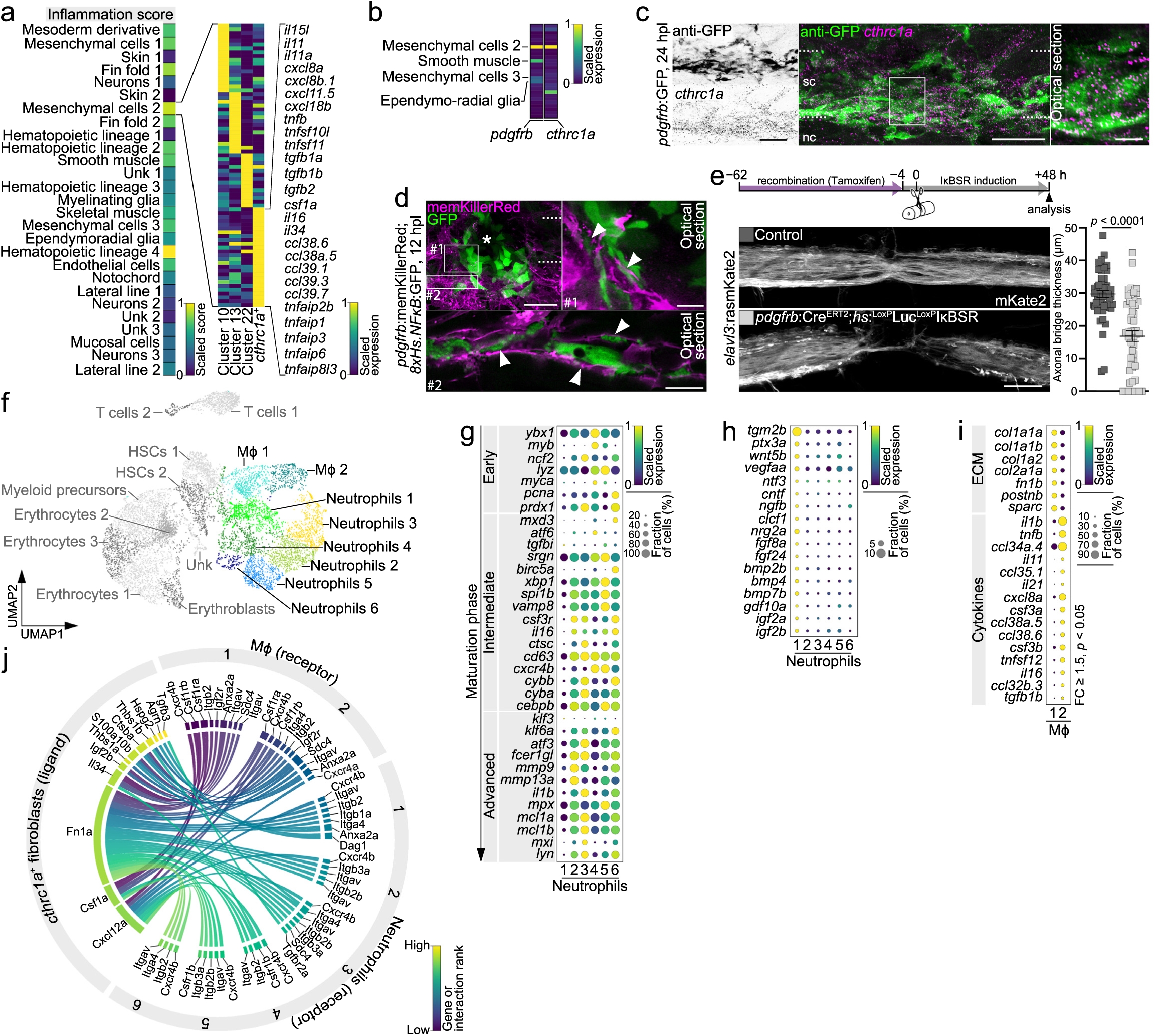
*cthrc1a*^+^ fibroblasts have an inflammatory profile. **a)** Scaled expression score of inflammation-related genes in major clusters and scaled expression of indicated genes in mesenchymal cells 2 subclusters. **b)** Scaled expression of *pdgfrb* and *cthrc1a* in all major clusters. Clusters with higher expression levels are indicated. **c)** *pdgfrb*^+^ cells (green) express *cthrc1a* mRNAs (magenta). *n* > 10. **d)** NFκB pathway reporter activity (cytoplasmic GFP; green) is detected in *pdgfrb*^−^ cells in the lesion core (asterisk) and in *pdgfrb*^+^ cells (membrane localized memKillerRed; magenta; arrowheads) at the wound margin. *n* > 10. **e)** *pdgfrb*^+^ cell-specific expression of IκB super repressor (IκBSR) after tamoxifen-induced Cre-mediated recombination leads to reduced axonal bridge thickness (white). Data are means ± SEM. Each data point represents one animal. Two-tailed Mann–Whitney test. *n*_control_ = 52, *n_pdgfrb_*_:IκBSR_ = 51 over three independent experiments. **f)** UMAP representation of all hematopoietic lineage clusters after second-level clustering. Mϕ and neutrophil clusters are highlighted in color. **g)** Scaled expression of genes associated with indicated maturation stages of neutrophils. **h)** Scaled expression of genes implicated in neuroprotection, neurogenesis, axon growth, and neurorepair pathways in neutrophil clusters. **i)** Scaled expression of cytokines and core matrisome genes (ECM) in Mϕ clusters. Shown are DEGs with FC ≥ 1.5 and *p* < 0.05. **j)** Predicted interactome of *cthrc1a^+^* fibroblast ligands with Mϕ and neutrophil receptors. Shown are 100 top ranked interactions. **a**–**j)** Images shown are maximum intensity projections of the lesion site (lateral view; rostral is left) or single optical sections (insets in **c**, **d**). Dashed lines indicate the spinal cord. Abbreviations: DEG, differentially expressed gene; ECM, extracellular matrix; h, hours; hpl, hours post-lesion; hs, hsp70l heat shock promoter; HSCs, hematopoietic stem cells; IκBSR, IκB super repressor; Mϕ, macrophages/monocytes; nc, notochord; sc, spinal cord; UMAP, uniform manifold approximation and projection; unk, unknown. Scale bars: 50 µm (**c**–**e**),10 µm (**c**, **d**; inset).

### Emergence of cthrc1a^+^ fibroblasts coincides with the innate immune response

To investigate whether *cthrc1a*^+^ fibroblasts interact with the immune response, we first performed second-level clustering of all four hematopoietic lineage clusters to identify effector cells of the innate immunity (Supplementary Fig. 3e). Based on commonly accepted markers, we identified two macrophage/monocyte (Mϕ) and six neutrophil transcriptional states (Fig. 3f; Supplementary Fig. 3f) ^73–78^. These Mϕ and neutrophil populations exhibited a heterogeneous cytokine expression profile, suggesting specialized roles in inflammation modulation during regeneration (Supplementary Fig. 3g). According to zebrafish-specific and pan-species marker genes, we categorized the neutrophil populations into different maturation phases (Fig. 3g) ^79^. Neutrophils 2 showed the highest expression of *mmp9* and *mmp13*, which typically indicates high maturity and phagocytic activity. Neutrophils 3 and 6 expressed intermediate levels of *mmp9* and high levels of *cybb* transcripts, reflecting advanced and intermediate maturation states. Neutrophils 4 and 5 expressed low *mmp9* and *mmp13* levels but high *cxcr4b* and *myca* levels, suggesting less mature states. The low expression of intermediate and advanced maturation markers in neutrophils 1 pointed to an early maturation stage. Notably, a fraction of neutrophils 1 expressed factors implicated in neuroprotection, neurogenesis, axon growth, and neurorepair pathways, including neurotrophins (*ntf3*, *ngfb*), neuropoietic cytokines (*cntf*, *clcf1*), neuregulins (*nrg2a*), and insulin-like growth factors (*igf2a/b*) (Fig. 3h). This is consistent with a previously described immature neutrophil population with regeneration-promoting functions in mammals ^80^. We did not detect clear differences between activation signatures of the two Mϕ populations, which supports the notion that macrophage polarization is a continuum rather than specific states ^81^. However, Mϕ 2 differentially expressed high levels of cytokines (e.g., *il1b*, *tnfb*, *cxcl8a*, *csf3a/b*), whereas Mϕ 1 exhibited high level expression of core matrisome genes (e.g., *col1a1a/b*, *col1a2*, *col2a1a*, *fn1b*) (Fig. 3i). Since macrophages have been reported to deposit ECM, including fibronectin and fibrillar collagens, in lesions of the spinal cord, kidney, and heart, Mϕ 1 likely represents a specific population involved in wound contraction and repair ^39,82–84^. Thus, our scRNA-seq approach captured the functional heterogeneity of Mϕ and neutrophils during spinal cord regeneration in zebrafish.

To assess the ability of *cthrc1a*^+^ fibroblasts to biochemically interact with the innate immune system, we predicted cellular interactions based on ligand and receptor gene expression. This analysis identified several potential ligand–receptor interactions between *cthrc1a*^+^ fibroblasts and different neutrophil/Mϕ populations, such as Csf1a–Csfr1a/b, Cxcl12a–Cxcr4b, and Tgfb3–Tgfbr2a (Fig. 3j). We then examined the temporal relationship between the appearance of *cthrc1a*^+^ fibroblasts and the innate immune response by comparing the relative cell number of the different cell populations during regeneration (Fig. 4a). This suggested that the appearance of the *cthrc1a*^+^ fibroblast state coincides with that of neutrophils 2 at 6 hpl. The peak number of *cthrc1a*^+^ fibroblasts was observed at 12 hpl, preceding the peak cell number of all other neutrophil populations and all Mϕ populations. In summary, *cthrc1a*^+^ fibroblasts are biochemically and temporally competent to interact with the innate immune response.

**Figure 4.**
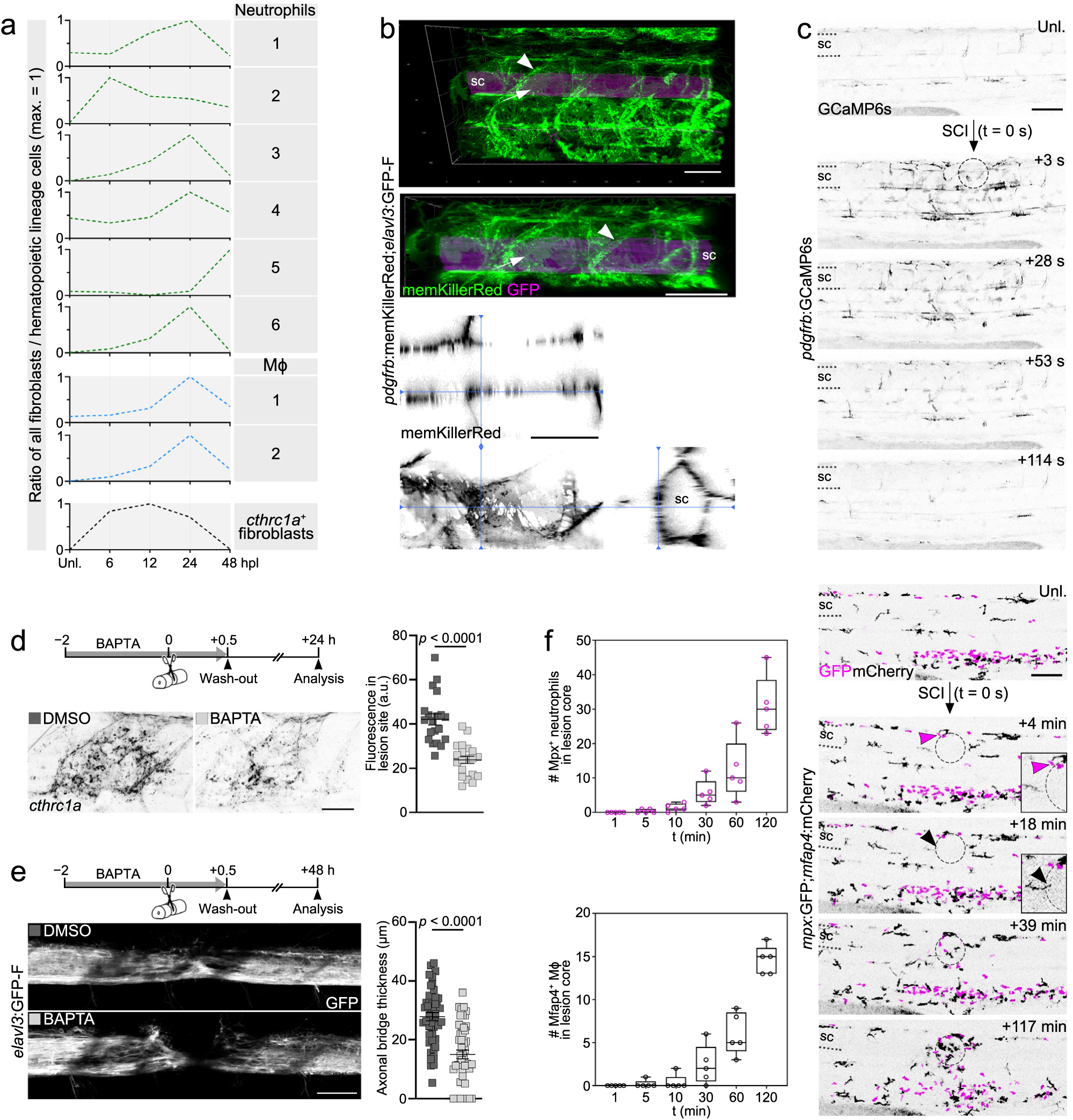
*cthrc1a*^+^ fibroblast activation precedes the innate immune response. **a)** Number of *cthrc1a*^+^ fibroblasts, Mϕ, and neutrophils in unlesioned animals and at indicated time points after SCI. Data are means normalized to the total number of fibroblasts or hematopoietic lineage cells, respectively. **b)** *pdgfrb*^+^ cells (memKillerRed; green/black) in the unlesioned zebrafish trunk. Arrowheads indicate *pdgfrb*^+^ cells in myotendinous junctions, arrows indicate *pdgfrb*^+^ cells covering the spinal cord (GFP; magenta). Images shown are 3D reconstructions and orthogonal projections (dorsal view, lateral view, and transversal view). *n* > 10. **c)** Time-lapse microscopy of *pdgfrb*:GCaMP6s transgenics reveals Ca^2+^ signals (black) in *pdgfrb*^+^ cells within 3 s after SCI. Images shown are single frames of Supplementary Video 1 (maximum intensity projections, lateral view; rostral is left). *n* = 5. **d and e)** Short-term treatment with the cell permeant Ca^2+^ chelator BAPTA-AM (abbreviated as BAPTA) leads to reduced *cthrc1a* mRNA expression in the lesion site and reduced axonal bridge thickness (white). Images shown are maximum intensity projections of the lesion site (lateral view; rostral is left). Two-tailed Student’s *t*-test. (**d**): *n* = 20 for each condition. (**e**): *n*_DMSO_ = 47, *n*_BAPTA_ = 48 animals over three independent experiments. **f)** Time-lapse microscopy reveals the dynamics of early Mpx^+^ neutrophil (magenta) and Mfap4^+^ Mϕ (black) recruitment after SCI. Arrowheads indicate the arrival of the first Mϕ and neutrophil at the lesion core. Images shown are single frames of Supplementary Video 2 (maximum intensity projections, lateral view; rostral is left). *n* = 5. **a**–**f)** Data are presented as mean ± SEM (**d**, **e**) or as box plots showing the median, first, and third quartiles with whiskers indicating the minimum and maximum values (**f**). Each data point represents one animal. Dashed lines indicate the spinal cord, dashed circle indicates lesion core (**c**, **f**). Abbreviations: a.u., arbitrary unit; hpl, hours post-lesion; max, maximum; min, minutes; Mϕ, macrophages/monocytes; s, seconds; sc, spinal cord; SCI, spinal cord injury; t, time; unl, unlesioned. Scale bars: 100 µm (**c**, **f**), 50 µm (**b**, **d**, **e**).

### Induction of the cthrc1a^+^ fibroblast state precedes the invasion of immune cells

Our scRNA-seq data indicated that neutrophil recruitment coincides with, or occurs after, the emergence of the *cthrc1a*^+^ fibroblast state. Based on this observation, we hypothesized that tissue-resident fibroblasts may play a role in sensing and mediating early wound detection. Consistent with this hypothesis, live imaging of mesenchymal cells 2 in *pdgfrb*:memKillerRed transgenic animals revealed an intricate fibroblast network that envelops the spinal cord and extends throughout the entire trunk in three dimensions (Fig. 4b). To investigate this further, we performed time-lapse imaging to visualize cytosolic calcium (Ca^2+^) spikes. Ca^2+^ influx through membrane channels and the release of Ca^2+^ from internal stores are among the earliest injury signals initiating tissue repair ^85^. To monitor Ca^2+^ spikes in mesenchymal cells 2, we used regulatory elements of the *pdgfrb* gene to drive expression of the GCaMP6s reporter (*pdgfrb*:GCaMP6s) ^86^. Analysis of time-lapse recordings showed a major burst of GCaMP6s activity in mesenchymal cells 2 within 5 s of SCI, which was detected up to 380 µm ± 102 µm rostral or caudal to the lesion center (Fig. 4c; Supplementary Video 1). GCaMP6s fluorescence returned to baseline levels after 171 s ± 20 s. This suggests rapid injury-associated changes in intracellular Ca^2+^ levels in mesenchymal cells 2. To probe the importance of early Ca^2+^ signaling on the *cthrc1a*^+^ fibroblast state, we used short-term treatments with either the cell-permeant Ca^2+^ chelator BAPTA-AM (abbreviated as BAPTA) or thapsigargin, which depletes Ca^2+^ in the endoplasmic reticulum, administered up to 30 min post-lesion ^87,88^. Both treatments led to reduced *cthrc1a* expression in the lesion site by 43% (BAPTA) and 78% (thapsigargin) at 24 hpl (Fig. 4d; Supplementary Fig. 4a). By contrast, constitutive *cthrc1a* expression in the inner ear and injury-induced expression of the matrisome genes *anxa2a* and *fn1b*, predominantly localized to other cell types, were unaffected (Supplementary Fig. 4b–f). Both short-term treatment paradigms also impaired axon regeneration: axonal bridge thickness was reduced by 46% and 53% in BAPTA- and thapsigargin-treated animals, respectively, compared to DMSO controls (Fig. 4e; Supplementary Fig. 4g). These findings suggest SCI triggers rapid Ca^2+^-mediated induction of the *cthrc1a*^+^ fibroblast state, a process which is required for effective axon regeneration. We next assessed the kinetics of neutrophil and Mϕ recruitment to the lesion core using time-lapse recordings and still images of *mpx*:GFP;*mfap4*:mCherry transgenic animals. The first neutrophil arrived in the lesion core 454 s ± 113 s post injury (Fig. 4f; Supplementary Video 2). Neutrophil numbers peaked between 4 and 6 hpl, consistent with previous reports (Supplementary Fig. 4h) ^15,17^. The earliest Mϕ was detected in the lesion core at 19 min ± 7 min post injury (Fig. 4f; Supplementary Video 2). Mϕ numbers peaked at 12 hpl and plateaued until 48 hpl (Supplemenatry Fig. 4h) ^15,17^. Altogether, these data suggest that wound detection by resident fibroblasts, which triggers the induction of the *cthrc1a*^+^ fibroblast state, precedes the recruitment of neutrophils and Mϕ following SCI.

### cthrc1a^+^ fibroblasts induce the innate immune response

Our data support that neutrophil and Mϕ recruitment occurs temporally after the activation of the *cthrc1a*^+^ fibroblast state, suggesting that these cells may play a role in inducing the innate immune response. To investigate this, we mitigated the emergence of *cthrc1a*^+^ fibroblasts using *pdgfrb*:NTR-mCherry transgenic animals, which enable targeted depletion of mesenchymal cells 2 via nitroreductase (NTR) activity upon exposure to metronidazole (MTZ) (Supplementary Fig. 5a). MTZ treatment efficiently ablated mCherry^+^ cells, although not all mesenchymal cells 2 were eliminated due to mosaic transgene expression (Supplementary Fig. 5a,b) ^13^. The induction of apoptosis in *pdgfrb*^+^ cells was confirmed using the TUNEL assay (Supplementary Fig. 5a). Importantly, the ablation of mesenchymal cells 2 in unlesioned animals did not lead to detectable recruitment of Mpx^+^ neutrophils to the fibroblast niche (Supplementary Fig. 5c). Following MTZ treatment, *cthrc1a* transcript levels in the lesion site were reduced by 80% at 24 hpl, indicating successful interference with the formation of *cthrc1a*^+^ fibroblasts (Supplementary Fig. 5d). This treatment regimen also led to the reduced appearance of Mpx^+^ neutrophils in the lesion core at 2 (−24%) and 6 hpl (−24%) and Mfap4^+^ Mϕ at 12 (−25%) and 24 hpl (−42%) (Supplementary Fig. 5e,f). To corroborate these findings, we employed *pdgfrb*:memKillerRed transgenic animals to induce localized cell death of mesenchymal cells 2 in the trunk prior to SCI (Fig. 5a; Supplementary Fig. 5g). KillerRed encodes a photosensitizer that generates reactive oxygen species upon light-induced photobleaching (referred to hereafter as irradiation), leading to site-specific oxidative stress and cell death ^89^. Four hours after irradiation, we confirmed *pdgfrb*^+^ cell-specific apoptosis using the TUNEL assay (Fig. 5a). At this time point, there was no detectable recruitment of Mpx^+^ neutrophils into the irradiated trunk (Supplementary Fig. 5h). Optogenetic local depletion of mesenchymal cells 2 led to reduced *cthrc1a* expression in the lesion site by 48% at 24 hpl, indicating impaired formation of *cthrc1a*^+^ fibroblasts (Fig. 5b). This treatment regimen also led to fewer Mpx^+^ neutrophils in the lesion core at 2 (−41%) and 6 hpl (−37%) and Mpeg1^+^ Mϕ at 12 (−37%) and 24 hpl (−25%) (Fig. 5c,d). Consistent with an attenuated immune response, axonal bridge thickness was reduced by 57% in irradiated *pdgfrb*:memKillerRed transgenic animals at 48 hpl (Supplementary Fig. 5i) ^15,19^. To independently confirm the results obtained from genetic manipulations, we applied the short-term BAPTA treatment paradigm to inhibit the emergence of *cthrc1a*^+^ fibroblasts. This intervention also led to fewer Mpx^+^ neutrophils in the lesion core at 2 (−29%) and 6 hpl (−29%) (Fig. 5e) and Mfap4^+^ Mϕ at 12 (−25%) and 24 hpl (−30%) (Fig. 5f). Importantly, bacterial lipopolysaccharide (LPS)-induced immune activation rescued the number of neutrophils and Mϕ in the lesion core in both optoablated (*pdgfrb*:memKillerRed) and BAPTA-treated animals (Fig. 5c–f) ^90^. This indicates that an alternative immunogenic signal can compensate for the absence of *cthrc1a*^+^ fibroblasts. Altogether, these data demonstrate that mitigating the induction of the *cthrc1a*^+^ fibroblast state attenuates the early innate immune response to SCI and inhibits axon regeneration.

**Figure 5.**
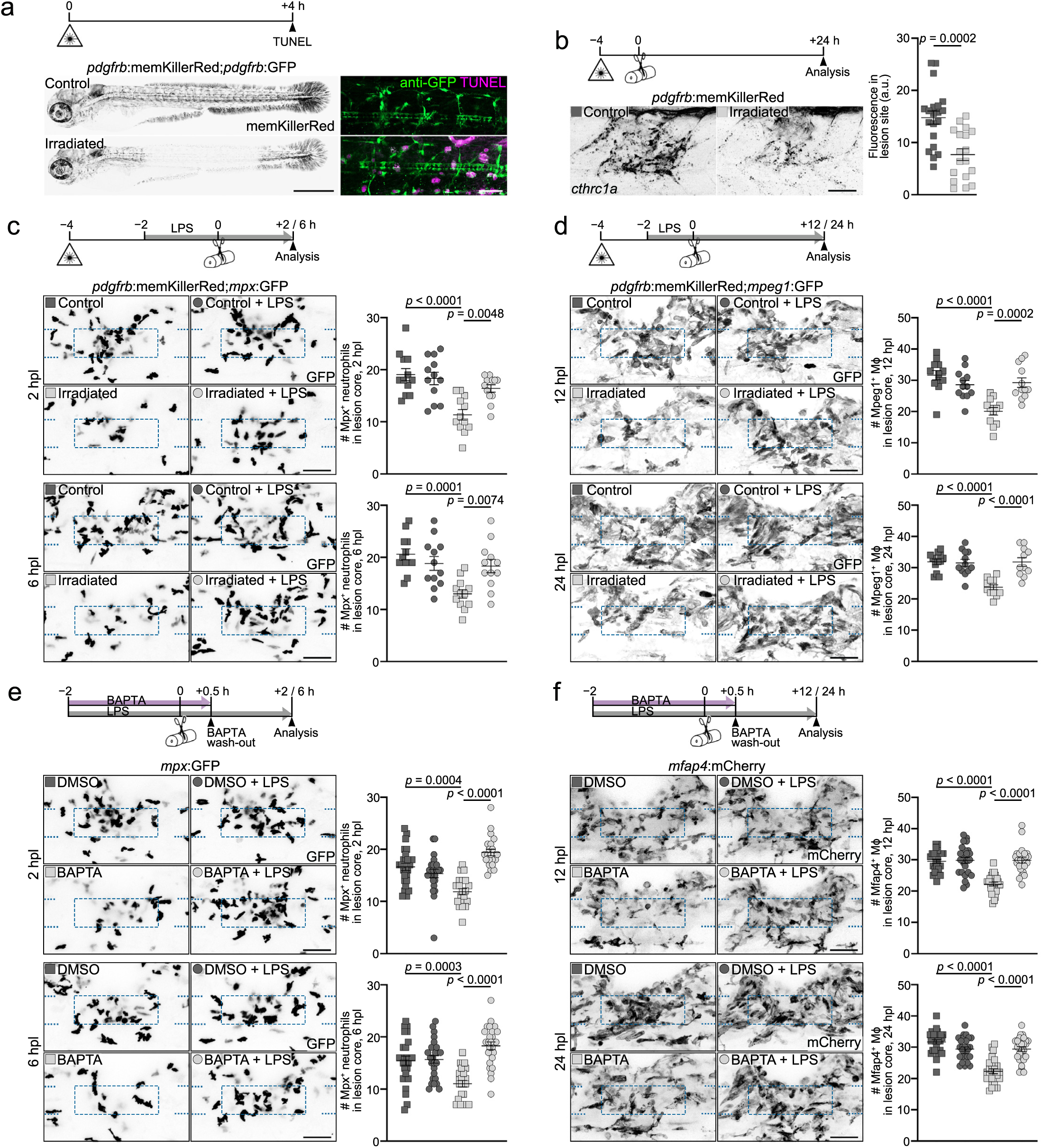
*cthrc1a*^+^ fibroblasts induce the inflammatory response after SCI. **a)** Exposure of *pdgfrb*:memKillerRed;*pdgfrb*:GFP transgenics to green light (hereafter referred to as irradiation) bleaches memKillerRed fluorescence (black) and induces apoptosis (TUNEL^+^; magenta) in *pdgfrb*^+^ cells (GFP; green). *n* = 15. **b)** Irradiation of *pdgfrb*:memKillerRed transgenics leads to reduced *cthrc1a* mRNA expression (black) in the lesion site. Two-tailed Student’s *t*-test. *n*_Control_ = 20, *n*_Irradiated_ = 18. **c and d)** Irradiation of *pdgfrb*:memKillerRed transgenics leads to a reduced appearance of Mpx^+^ neutrophils (black) (**c**) and Mpeg1^+^ Mϕ (black) (**d**) in the lesion core. Treatment with LPS rescues neutrophil and Mϕ numbers in irradiated animals. One-way Anova with Šídák’s multiple comparisons test. Neutrophils: *n* = 12 for each condition. Mϕ(12 hpl): *n* = 12 for each condition. Mϕ(24 hpl): *n*_Control_ = 12, *n*_LPS_ = 12, *n*_Irradiated_ = 12, *n*Irradiated+LPS = 11. **e and f)** Short-term treatment with the cell permeant Ca^2+^ chelator BAPTA-AM (BAPTA) leads to a reduced appearance of Mpx^+^ neutrophils (black) (**e**) and Mfap4^+^ Mϕ (black) (**f**) in the lesion core. LPS treatment rescues neutrophil and Mϕ numbers in BAPTA-treated animals. Neutrophils(2 hpl): Kruskal–Wallis test; *n*_DMSO_ = 25, *n*_DMSO+LPS_ = 23, *n*_BAPTA_ = 21, *n*_BAPTA+LPS_ = 22. Neutrophils(6 hpl): One-way Anova with Šídák’s multiple comparisons test; *n*_DMSO_ = 26, *n*_DMSO+LPS_ = 26, *n*_BAPTA_ = 25, *n*_BAPTA+LPS_ = 26. One-way Anova with Šídák’s multiple comparisons test. Mϕ(12 hpl): *n*_DMSO_ = 25, *n*_DMSO+LPS_ = 24, *n*_BAPTA_ = 25, *n*_BAPTA+LPS_ = 24. Mϕ(24 hpl): *n* = 27 for each condition. **a**–**f)** Data are means ± SEM. Each data point represents one animal. Images shown are maximum intensity projections of uninjured animals, uninjured trunk, or lesion site (lateral view; rostral is left). Dashed lines indicate spinal cord, dashed square indicates region of quantification. Scale bars: 500 µm (overview in **a**) and 50 µm (**a**–**f**). Abbreviations: a.u., arbitrary unit; h, hours; hpl, hours post-lesion; LPS, bacterial lipopolysaccharide; Mϕ, macrophages/monocytes; SCI, spinal cord injury.

### cthrc1a^+^ fibroblasts control inflammation resolution

To further characterize the dynamics of inflammation in *cthrc1a*^+^ fibroblast-depleted animals, we performed scRNA-seq analysis of *pdgfrb*:NTR-mCherry transgenic animals treated with either MTZ or DMSO control at 24 hpl, after the acute inflammatory phase (Fig. 6a). Following quality control, 36,450 cells across two biological replicates per condition were analyzed. Second-level clustering of the hematopoietic lineage cells identified two Mϕ and three neutrophil populations (Supplementary Fig. 6a–d). The transcriptional states of the three neutrophil populations corresponded to early (neutrophils 1), intermediate (neutrophils 2), and advanced (neutrophils 3) maturation stages (Fig. 6b,c) ^79^. Consistent with the first scRNA-seq dataset, early maturation stage neutrophils 1 expressed genes associated with neurorepair pathways, suggesting a regeneration-promoting role (Fig. 6d) ^80^. Comparison of relative cell numbers between conditions revealed a shift toward intermediate and advanced neutrophil maturation stages in *cthrc1a*^+^ fibroblast-depleted animals (Fig. 6b). Specifically, the number of neutrophils 1 (pro-neurorepair profile) was reduced in MTZ-treated *pdgfrb*:NTR-mCherry transgenics while neutrophils 2 and 3 (pro-inflammatory profiles) expanded compared to DMSO-treated controls. Additionally, neutrophils 2 and 3 in MTZ-treated *pdgfrb*:NTR-mCherry transgenics, showed increased expression of pro-inflammatory cytokines, including *il1b*, *il4*, *il16*, *il21*, *il34*, *tnfb*, and *tnfsf18* (Fig. 6d). In contrast, neutrophils 1 exhibited reduced expression of genes involved in neuroprotection, neurogenesis, axon growth, such as *wntb5b*, *vegfaa*, *ntf3*, *ngfa*, *nrg1*, *nrg2a*, *fgf8a*, *bmp1b*, *bmp6*, *gdf6a*, and *igf2b* (Fig. 6d). These findings indicate a shift in neutrophil profiles toward a pro-inflammatory phenotype in *cthrc1a*^+^ fibroblast-depleted animals. Pseudotime analysis supported a transcriptional transition in neutrophils between DMSO and MTZ-treated conditions (Supplementary Fig. 6e). We next analyzed the two Mϕ populations (Fig. 6e). Differential expression analysis identified Mϕ 1 as ECM-producing repair macrophages, while Mϕ 2 displayed elevated cytokine expression (Fig. 6f). Similar to neutrophils, the relative cell number of Mϕ 1 was reduced in *cthrc1a*^+^ fibroblast-depleted animals, while Mϕ 2 increased (Fig. 6e). Furthermore, Mϕ 2 in MTZ-treated *pdgfrb*:NTR-mCherry transgenic animals exhibited higher expression levels of pro-inflammatory cytokines, including *il21*, *il34*, *tnfb*, *cxcl8b.1*, *tnfsf11*, *tnfsf12*, *ccl34a.4*, *ccl35.1*, and *ccl38* (Fig. 6g). Pseudotime analysis reflected this transitioning of Mϕ transcriptional states between DMSO and MTZ-treated conditions (Supplementary Fig. 6f). These findings indicate that mitigating the induction of the *cthrc1a*^+^ fibroblast state leads to an exacerbated and prolonged inflammatory response. To experimentally validate the role of *cthrc1a*^+^ fibroblasts in curtailing inflammation, we optogenetically depleted *cthrc1a*^+^ fibroblasts in the lesion site in *pdgfrb*:memKillerRed transgenic animals at 6 hpl, corresponding to the peak of neutrophil influx. This resulted in a 3.5-fold increase in neutrophil numbers in the lesion core at 24 hpl compared to controls (Fig. 6h). The depletion of *cthrc1a*^+^ fibroblasts at the peak of inflammation therefore leads to the retention of neutrophils in the lesion site. Altogether, these data indicate that *cthrc1a*^+^ fibroblasts regulate the resolution of neutrophil-driven inflammation after SCI.

**Figure 6.**
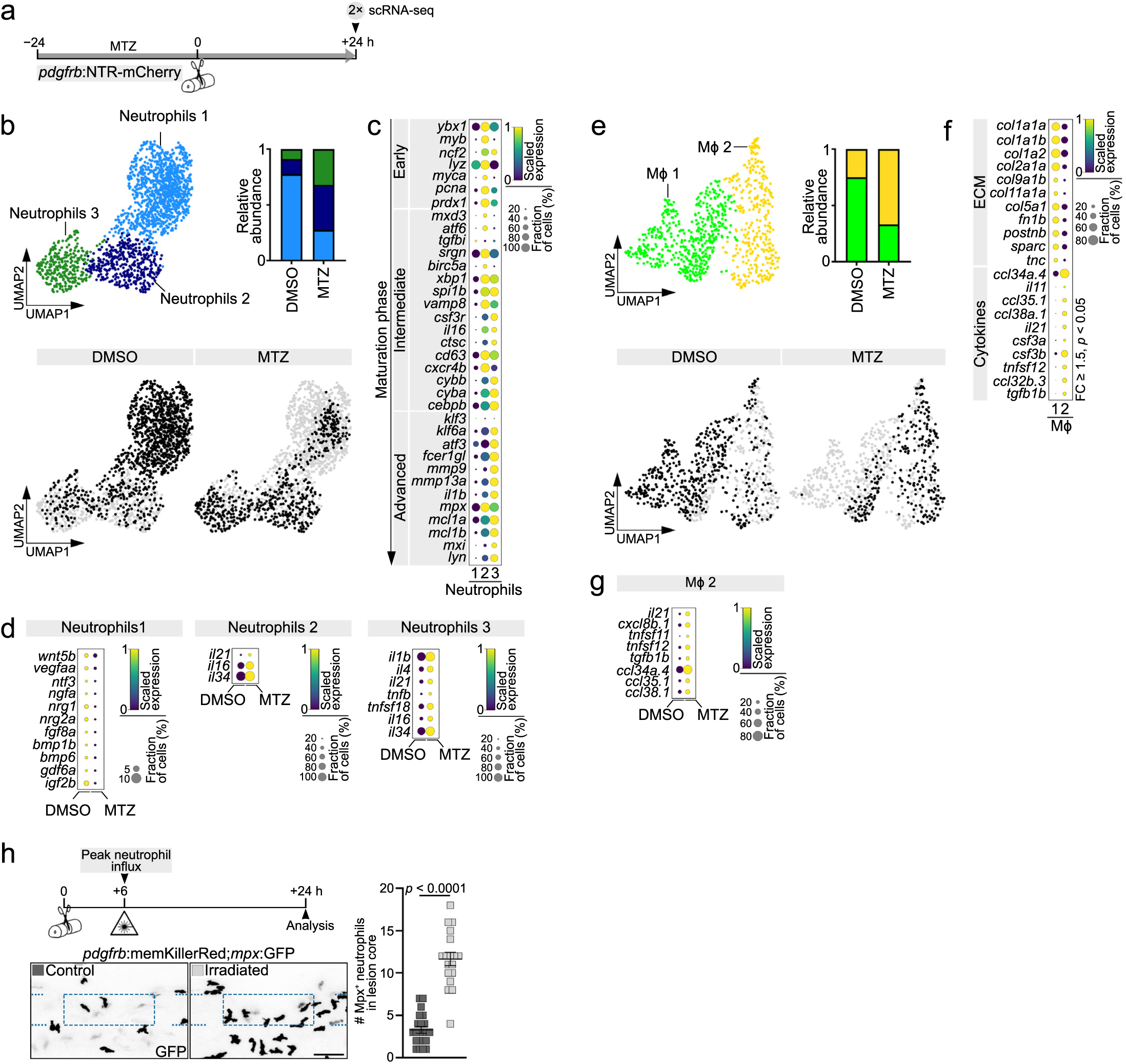
*cthrc1a*^+^ fibroblasts control inflammation resolution. **a)** Timeline of the scRNA-seq experiment shown in **b**–**g**. Cells for scRNA-seq analysis were isolated from DMSO or MTZ-treated *pdgfrb*:NTR-mCherry transgenics. **b)** Shown are UMAP representations of all neutrophils after second-level clustering and relative abundance of the subclusters in each condition (DMSO/MTZ). **c)** Scaled expression of genes associated with indicated maturation stages of neutrophils. **d)** Scaled expression of indicated genes in neutrophil subclusters in each condition (DMSO/MTZ). For neutrophils 1, factors implicated in neuroprotection, neurogenesis, axon growth, and neurorepair pathways are plotted. For neutrophils 2 and 3, pro-inflammatory cytokines are plotted. **e)** Shown are UMAP representations of all Mϕ after second-level clustering and relative abundance of the subclusters in each condition (DMSO/MTZ). **f)** Scaled expression of indicated core matrisome genes (ECM) and cytokines in Mϕ clusters. Shown are DEGs with FC ≥ 1.5 and *p* < 0.05. **g)** Scaled expression of indicated pro-inflammatory cytokines in Mϕ cluster 2 in each condition (DMSO/MTZ). **h)** Exposure of *pdgfrb*:memKillerRed transgenics to green light at 6 hpl leads to an increased number of Mpx^+^ neutrophils (black) in the lesion core. Images shown are maximum intensity projections of the lesion site (lateral view; rostral is left). Dashed lines indicate the spinal cord, dashed square indicates the region of quantification. Data are means ± SEM. Each data point represents one animal. Two-tailed Mann–Whitney test. *n*_Control_ = 22, *n*_Irradiated_ = 18 over two independent experiments. Scale bar: 50 µm. **a**–**h)** Abbreviations: h, hours; hpl, hours post-lesion; Mϕ, macrophages/monocytes; MTZ, metronidazole; UMAP, uniform manifold approximation and projection.

### cthrc1a^+^ fibroblasts control inflammation through cytokine release

To identify mediators of the inflammation-modulating functions on neutrophils, we screened for transcripts of genes encoding secreted inflammation-associated factors in *cthrc1a*^+^ fibroblasts (Supplementary Fig. 7a). Among these, *cxcl8b.1*, encoding the chemoattractant Cxcl8 (IL-8), which binds to the Cxcr1/2 receptor on neutrophils, was upregulated in mesenchymal cells 2 after SCI and peaked in *cthrc1a*^+^ fibroblasts at 6 hpl (Fig. 7a,b) ^17,91^. Comparing *cxcl8b.1* transcript levels across all major clusters in the scRNA-seq dataset revealed mesenchymal cells 2 as the principal source, while hematopoietic lineage cells showed only low-level expression (Fig. 7c; Supplementary Fig. 7b). ISH confirmed induction of *cxcl8b.1* expression in *pdgfrb^+^* mesenchymal cells 2 at the wound margin as early as 3 hpl (Fig. 7d; Supplementary Fig. 7c,d). To test the relevance of *cthrc1a*^+^ fibroblast-derived Cxcl8b.1 for neutrophil recruitment, we used an established splice-blocking morpholino targeting the *cxcl8b.1* pre-mRNA (*cxcl8b.1* splice MO) ^91,92^. To disrupt the SCI-induced synthesis of Cxcl8b.1 proteins and to account for proteins that are synthesized prior to injury for immediate release, we delivered the *cxcl8b.1* splice MO into the zygote. Knockdown of *cxcl8b.1* reduced the number of Mpx^+^ neutrophils in the lesion core at 2 (−23%) and 6 hpl (−19%) (Fig. 7e). Similarly, injection of a translation-blocking antisense morpholino targeting *cxcl8b.1* (*cxcl8b.1* ATG MO) led to a reduced influx of Mpx^+^ neutrophils at 2 hpl (−36%) (Supplementary Fig. 7e). To corroborate these results, we inhibited Cxcr1/2 signaling using the compound SB225002, which resulted in fewer Mpx^+^ neutrophils in the lesion core at 2 (−73%) and 6 hpl (−26%) (Fig. 7f) ^17^. Importantly, axon regeneration was reduced by 41% in *cxcl8b.1* splice MO-treated animals (Fig. 7g). These data indicate that Cxcl8b.1 partly mediates the inflammation inducing role of *cthrc1a*^+^ fibroblasts on neutrophils.

**Figure 7.**
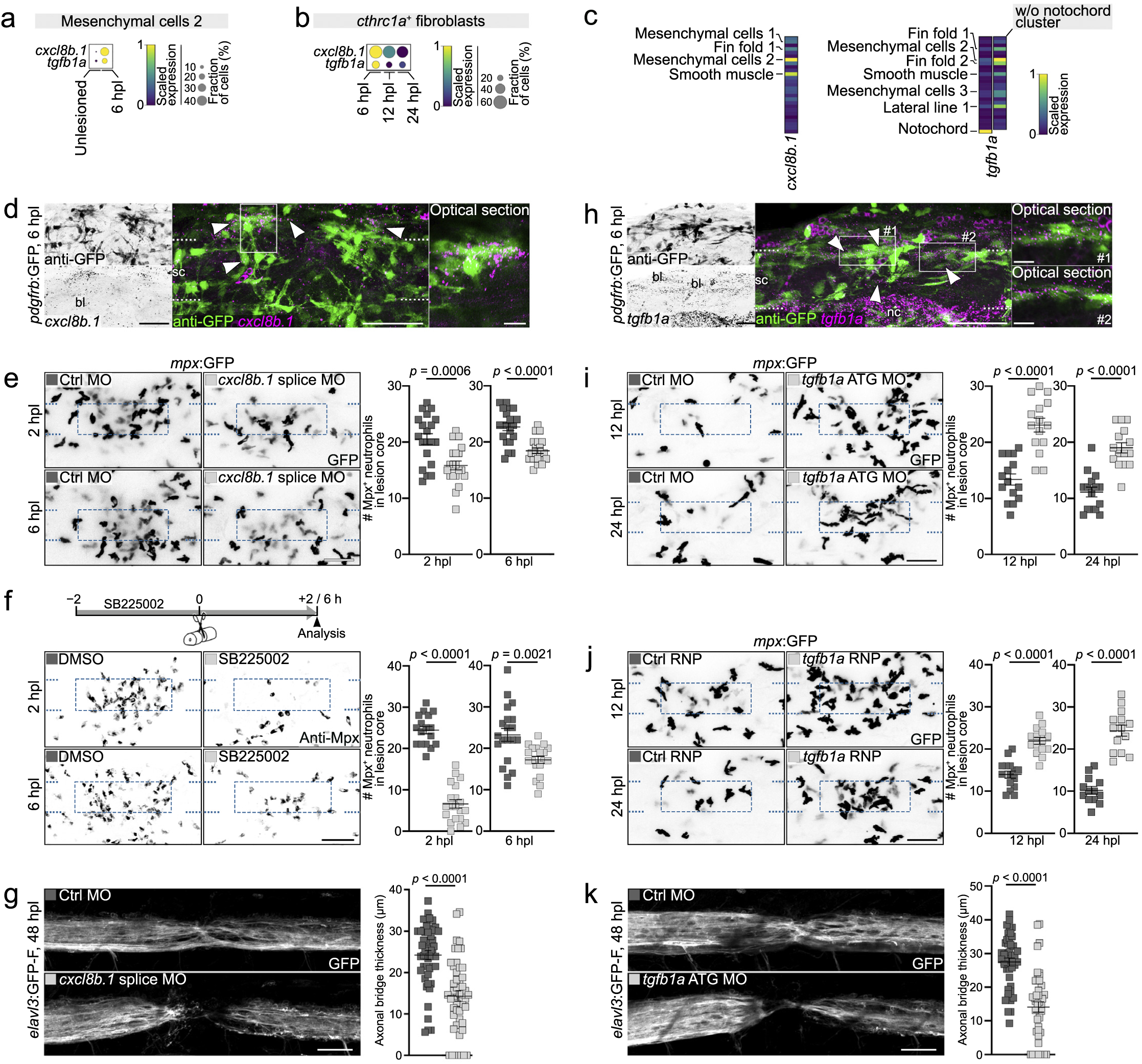
Cxcl8 and Tgfβ1 mediate the inflammation-modulating functions of *cthrc1a*^+^ fibroblasts. **a and b)** Scaled expression of *cxcl8b.1* and *tgfb1a* in mesenchymal cells 2 (**a**) and *cthrc1a*^+^ fibroblasts (**b**) at indicated time points. **c)** Scaled expression of *cxcl8b.1* and *tgfb1a* in all major clusters. Clusters with high expression levels are indicated. Note: *cxcl8b.1* transcript abundance is highest and *tgfb1a* transcript abundance is second highest in mesenchymal cells 2. **d)** *pdgfrb*^+^ cells (green) express *cxcl8b.1* mRNA (magenta; arrowheads). *n* > 10. **e)** Delivery of a *cxcl8b.1* splice blocking morpholino (splice MO) leads to reduced appearance of Mpx^+^ neutrophils (black) in the lesion core. Two-tailed Student’s *t*-test. *n* = 20 for each condition. **f)** Treatment with the Cxcr1/2 antagonist SB225002 leads to reduced appearance of Mpx^+^ neutrophils (black) in the lesion core. Two-tailed Student’s *t*-test. 2 hpl: *n*_DMSO_ = 17, *n*SB225002 = 20. 6 hpl: *n*DMSO = 22, *n*SB225002 = 21. **g)** Delivery of a *cxcl8b.1* splice MO leads to reduced axonal bridge thickness (white). Two-tailed Student’s *t*-test. *n*_Ctrl MO_ = 50, *n_cxcl8b.1_*_splice MO_ = 49 over three independent experiments. **h)** *pdgfrb*^+^ cells (green) expres *tgfb1a* mRNA (magenta; arrowheads). *n* > 10. **i)** Delivery of a *tgfb1a* translation blocking morpholino (ATG MO) leads to increased retention of Mpx^+^ neutrophils (black) in the lesion core. 12 hpl: Two-tailed Student’s *t*-test, *n* = 15 for each condition. 24 hpl: Two-tailed Mann–Whitney test, *n* = 15 for each condition. **j)** CRISPR manipulation of *tgfb1a* leads to increased retention of Mpx^+^ neutrophils (black) in the lesion core. 12 hpl: Two-tailed Student’s *t*-test, *n* = 15 for each condition. 24 hpl: Two-tailed Student’s *t*-test, *n*_Ctrl RNP_ = 13, *n_tgfb1a_* _RNP_ = 14. **k)** Delivery of a *tgfb1a* ATG MO leads to reduced axonal bridge thickness (white). Two-tailed Student’s *t*-test. *n* = 45 for each condition over three independent experiments. **a**–**k)** Data are means ± SEM. Each data point represents one animal. Images shown are maximum intensity projections of the lesion site (lateral view; rostral is left) or single optical sections (insets). Dashed lines indicate the spinal cord, dashed square indicates the region of quantification, solid square indicates the location of the inset. Scale bars: 50 µm (**d**–**k**), 10 µm (insets in **d**, **h**). Abbreviations: bl, blood; ctrl, control; h, hours; hpl, hours post-lesion; MO, morpholino; Mϕ, macrophages/monocytes; nc, notochord; RNP, gRNA/Cas9 ribonucleoprotein complex; sc, spinal cord; w/o, without.

To identify mediators of the inflammation-resolving function of *cthrc1a*^+^ fibroblasts, we focused on the immunosuppressive Tgfβ cytokine family member Tgfβ1a, whose expression was upregulated in mesenchymal cells 2 after SCI (Fig. 7a). Across all major clusters, *tgfb1a* transcripts were predominantly detected in different fibroblast-like cell populations and exhibited the second-highest expression in mesenchymal cells 2, while hematopoietic lineage cells expressed negligible levels of *tgfb1a* (Fig. 7c; Supplementary Fig. 7f). In *cthrc1a*^+^ fibroblasts, *tgfb1a* expression peaked when neutrophil influx reached its maximum at 6 hpl (Fig. 7b). ISH confirmed the induction of *tgfb1a* transcripts in *pdgfrb^+^*mesenchymal cells 2 at the wound margin at 6 hpl (Fig. 7h). Injection of a *tgfb1a* ATG MO into the zygote led to a 1.7-fold increase in the number of Mpx^+^ neutrophils in the lesion core at 12 and 24 hpl (Fig. 7i) ^93^. To corroborate these findings, we injected gRNAs targeting the *tgfb1a* gene together with Cas9 protein (*tgfb1a* RNP) into the zygote for efficient gene disruption, which also resulted in an increase in neutrophil numbers in the lesion core at 12 and 24 hpl (1.6- and 2.4-fold increase, respectively) (Fig. 7j; Supplementary Fig. 7g) ^20^. Thus, knockdown of *tgfb1a* retains neutrophils in the lesion core, delaying inflammation resolution. Consistent with a prolonged inflammatory phase, axon regeneration was reduced by 49% in *tgfb1a* ATG MO-injected animals at 48 hpl (Fig. 7k) ^15^. These data indicate that Tgfβ1a partly mediates the inflammation-resolving function of *cthrc1a*^+^ fibroblasts on neutrophils.

Altogether, our data support that *cthrc1a*^+^ fibroblasts biphasically control inflammation through the release of immune-modulating cytokines.

### Disturbing inflammation dynamics alters the mechano-structural tissue properties

In *cthrc1a*^+^ fibroblast-depleted animals, axon regeneration is impaired in part because of aberrant inflammation dynamics. To investigate the impact of an altered inflammatory phase on the lesion environment’s mechanical and structural properties, we treated zebrafish with the Cxcr1/2 antagonist SB225002 (Fig. 8a; Supplementary Fig. 8a) ^13,14^. This treatment mimicked the effect of *cthrc1a*^+^ fibroblast depletion on neutrophils but did not affect *cthrc1a* expression or Mfap4^+^ Mϕ recruitment to the lesion core (Fig. 7f; Fig. 8b,c; Supplementary Fig. 8b) ^17^. Specifically, at 2 and 6 hpl, fewer Mpx^+^ neutrophils were recruited to the lesion core, whereas a higher number of neutrophils were retained at 12, 24, and 48 hpl (1.4-, 2.3-, and 2.4-fold increase, respectively). Moreover, SB225002 treatment inhibited axon regeneration (−86%) and recovery of swimming distance (−70%) compared to DMSO controls (Fig. 8d,e; Supplementary Fig. 8d). Thus, pharmacological interference with Cxcr1/2 signaling allows us to elucidate the impact of an altered neutrophil response on the lesion environment’s properties.

**Figure 8.**
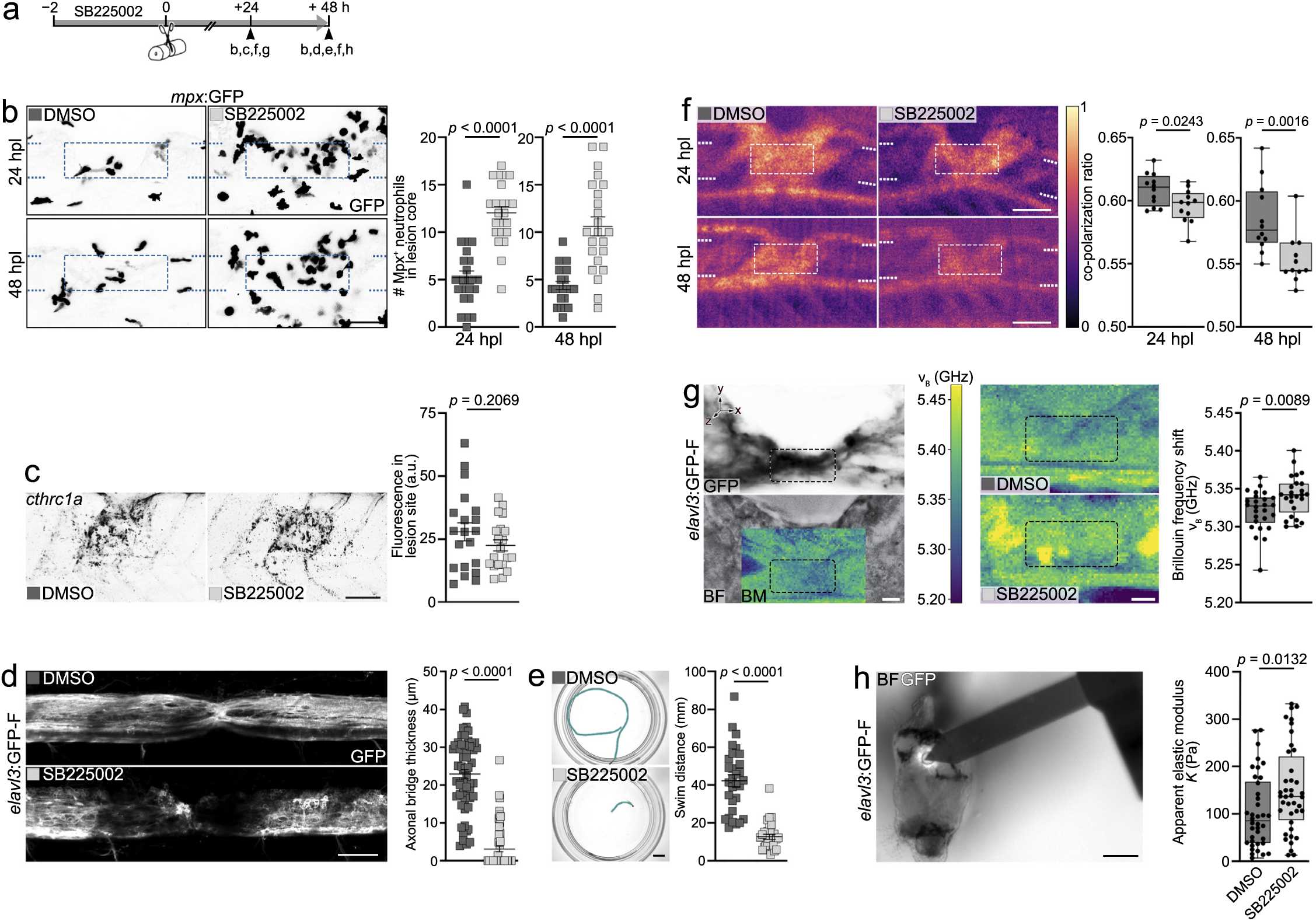
Disturbing inflammation dynamics alters the mechano-structural properties of the lesion environment. **a)** Timeline for treatments with the Cxcr1/2 antagonist SB225002 shown in **b**–**h**. **b)** SB225002 treatment leads to increased retention of Mpx^+^ neutrophils (black) in the lesion core. Two-tailed Mann–Whitney test, *n* = 25 for each condition. **c)** SB225002 treatment does not reduce *cthrc1a* mRNA expression (black) in the lesion site. Two-tailed Student’s *t*-test, *n* = 20 for each condition. **d)** SB225002 treatment leads to reduced axonal bridge thickness (white). Two-tailed Mann– Whitney test, *n* = 61 for each condition over three independent experiments. **e)** SB225002 treatment impairs recovery of swimming distance. Two-tailed Mann–Whitney test, *n* = 30 for each condition. **f)** SB225002 treatment decreases the co-polarization ratio of the lesion site. 24 hpl: Two-tailed Student’s *t*-test, *n* = 12 for each condition. 48 hpl: Two-tailed Mann–Whitney test, *n*DMSO = 12, *n*SB225002 = 11. **g)** SB225002 treatment increases *v*_B_ of the lesion core. Images shown are brightfield intensity, confocal fluorescence, or Brillouin frequency shift map through the center of the lesion site. Two-tailed Student’s *t*-test, *n*_DMSO_ = 26, *n*_SB225002_ = 25 over six independent experiments. **h)** SB225002 treatment increases *K* of the lesion core. Two-tailed Mann–Whitney test, *n*_DMSO_ = 38, *n*_SB225002_ = 40 over six independent experiments. **a**–**h)** Data are presented as mean ± SEM (**b**–**e**) or as box plots showing the median, first, and third quartiles with whiskers indicating the minimum and maximum values (**f**–**h**). Each data point represents one animal. Images shown are maximum intensity projections (**b**–**d**; lateral view; rostral is left), average intensity projections (**f**; lateral view; rostral is left), sagittal optical sections (**g**; lateral view; rostral is left), or transversal view of the lesion site (dorsal is up). Dashed lines indicate the spinal cord, dashed square indicates the region of quantification. Scale bars: 5 mm (**e**), 100 µm (**h**), 50 µm (**b**–**d**, **f**), 10 µm (**g**). Abbreviations: a.u., arbitrary unit; BF, brightfield intensity; BM, Brillouin frequency shift map; h, hours; hpl, hours post-lesion.

To assess structural changes, we applied *in vivo* cross-polarized optical coherence tomography (CP-OCT). CP-OCT measures relative changes in the polarization of incident light, offering enhanced contrast to highlight birefringent properties of the tissue ^11,94,95^. The co-polarization ratio—the ratio of preserved polarization to total reflectivity—was reduced in the lesion site in SB225002-treated animals at 24 (DMSO: 0.609 ± 0.004, SB225002: 0.596 ± 0.004) and 48 hpl (DMSO: 0.586 ± 0.008, SB225002: 0.554 ± 0.006) compared to DMSO controls (Fig. 8f). These findings suggest that the structural properties of the lesion environment change when inflammation dynamics are aberrant.

To assess mechanical properties of the lesion environment *in vivo*, we first used Brillouin microscopy (BM) ^11,96–98^. BM quantifies the Brillouin frequency shift (ν_B_) at GHz range, an inelastic scattering process by the interaction between light and thermally induced density fluctuations in the sample. ν_B_ is dependent on the longitudinal modulus (*M*’ with ν_B_*∝* √*M* ′), which describes the inverse of the longitudinal compressibility, on the density of the tissue, and on its refractive index (*n*). In SB225002-treated animals, we observed a mean increase in ν_B_ of 20 MHz in the lesion core at 24 hpl compared to controls, indicating a decreased compressibility (DMSO: 5.321 GHz ± 0.005 GHz, SB225002: 5.341 GHz ± 0.005 GHz) (Fig. 8g). We did not detect significant differences in the refractive index distribution of the lesion core using *in vivo* optical diffraction tomography, supporting that the observed changes in ν_B_ are primarily due to changes in *M*’ of the sample (Supplementary Fig. 8e) ^11,99^ . We next used atomic force microscopy (AFM)-based indentation measurements to quantify the reduced apparent elastic modulus (*K*) —a measure of tissue stiffness—of the lesion core in living tissue preparations. In SB225002-treated animals, we observed a 1.5-fold increase in *K* at 48 hpl, indicating a stiffening of the tissue (Fig. 8h). Thus, AFM and BM indicate changes in the mechanical response of the lesion environment at different time and length scales when neutrophil dynamics are aberrant.

Collectively, these data indicate that disturbing inflammation dynamics alters the mechano-structural properties of the lesion environment and inhibits axon regeneration after SCI.

## DISCUSSION

Axon regeneration in the mammalian CNS is inhibited by chronic inflammation and fibrous scarring, driven by the complex interactions between fibroblasts and immune cells ^2–6^. In contrast, we demonstrate that in zebrafish, the interplay between fibroblasts and innate immune cells facilitates regeneration. Our findings support a model in which tissue-resident fibroblasts rapidly detect SCI and adopt a transient, less differentiated, inflammation-related, non-fibrotic state. During the early acute phase of injury, these reactive fibroblasts secrete pro-inflammatory cytokines to induce and guide the innate immune response, a crucial step for initiating CNS regeneration ^15,17,100^. As neutrophil influx peaks in the late acute phase, fibroblasts undergo a functional shift, producing immunosuppressive cytokines to resolve inflammation, which is essential for axon regeneration in the subacute phase of injury. This biphasic regulation limits the neutrophil-driven acute inflammatory phase, preventing chronic inflammation, thereby fostering a lesion environment conducive to regeneration. Experimentally altering inflammation dynamics leads to changes in the mechano-structural properties of the lesion environment, which may stem from disrupted neutrophil functions, including debris clearance, secretion of ECM-modulating enzymes, and modulation of fibroblast activity and ECM secretion ^101^. Changes in mechanical tissue properties, such as elasticity and viscosity, have been shown to correlate with neural regeneration, offering a possible explanation for the reduced regenerative capacity of animals with altered inflammation dynamics ^7,9–11,102,103^. Furthermore, our study highlights Cxcl8 and Tgfβ1 as cytokines that mediate, at least in part, the dual role of fibroblasts in both inducing and resolving inflammation.

The injury-specific fibroblast population identified in this study is defined by high expression levels of *cthrc1a*, which encodes a glycoprotein critical for spinal cord regeneration in zebrafish ^13^. Interestingly, *CTHRC1* has also been recognized as a marker of a unique fibroblast population in fibrotic conditions in mammals, including humans ^104,105^. While *CTHRC1^+^* fibroblasts in both mammals and zebrafish share an inflammation-related profile, they exhibit significant differences in differentiation status and associated ECM secretion ^13,104,105^. In mammals, *CTHRC1^+^* fibroblasts produce high levels of ECM proteins, whereas their zebrafish counterparts display attenuated expression of matrisome genes, including those encoding known inhibitors of axon regeneration. We propose that zebrafish *cthrc1a^+^* fibroblasts emerge through dedifferentiation, a process linked to reduced ECM secretion ^106^. Notably, fibroblast dedifferentiation has also been proposed as a critical mechanism contributing to organ regeneration in other vertebrate species ^107,108^. Thus, two major processes that hinder axon regeneration in the mammalian CNS—persistent inflammation and fibroblast-derived ECM deposition—are coordinated and fine-tuned in zebrafish to promote regeneration. The present study therefore underscores a fundamental distinction in the fibroblast response to SCI and its impact on regeneration outcomes between zebrafish and mammals, including humans.

Neutrophils are traditionally regarded as the first responders to injury, with their recruitment to distant wounds guided by tissue-resident macrophages ^109^. Various mechanisms have been implicated in the early detection of wounds by immune cells, including reactive oxygen species, osmotic and ion gradients, the release of nucleotides, and Ca^2+^ waves emanating from the site of injury ^110–114^. Here, we present evidence that changes in intracellular Ca^2+^ levels in fibroblasts precede and facilitate the recruitment of neutrophils and macrophages. Our data supports the notion that fibroblasts form a body-wide cellular network, which is capable of sensing mechanical force-damage, possibly involving mechanosensitive ion channels such as Piezo ^115,116^. We therefore propose that fibroblasts function as the primary wound-detecting cells, acting upstream of the innate immune response. This model is supported by our observations that mitigating the emergence of *cthrc1a^+^* fibroblasts reduces the recruitment of neutrophils and macrophages to the lesion core, an effect that can be rescued by LPS treatment. Moreover, in larval zebrafish, tissue-resident macrophages are present at much lower density than fibroblasts, making them unlikely to serve as primary cells guiding neutrophil recruitment ^117^. The detection of injury-induced GCaMP reporter activity in fibroblasts several hundred micrometer away from the lesion center provides further evidence for a model in which fibroblasts propagate injury signals through gap junction-mediated direct cell–cell communication, mediating the detection of distant wounds by neutrophils.

In conclusion, our findings establish the biphasic control of inflammation by dedifferentiated fibroblasts as a key mechanism for CNS regeneration. Thus, identifying the cues that govern fibroblast dedifferentiation and inflammation control in zebrafish has potential implications for CNS injury in mammals, where regeneration fails, partly because of sustained inflammation and fibrosis.

## METHODS

### Zebrafish husbandry and transgenic lines

Zebrafish lines were kept and reared at the Max Planck Institute for the Science of Light and the Max-Planck-Zentrum für Physik und Medizin under a 14/10 h light-dark cycle at 28.5°C and according to FELASA recommendations ^118^, and the supervision of the veterinary authorities of the city Erlangen (permits: I/39/FNC and VII/390/DL003 of the Amt für Veterinärwesen und gesundheitlichen Verbraucherschutz Stadt Erlangen). All experimental procedures were done on larval zebrafish aged up to 120 hours post-fertilization (hpf), derived from voluntarily mating adult zebrafish. Procedures on zebrafish larvae up to 120 hpf are not regulated as animal experiments by the European Commission Directive 2010/63/EU.

We used AB and WIK wild-type strains of zebrafish (*Danio rerio*) and the following transgenic zebrafish lines: BAC(*pdgfrb*:Gal4ff)^ncv24^ ^119^, *UAS*:EGFP^zf82^ ^120^, *UAS-E1b*:Eco.NfsB-mCherry^c264 121^, *pdgfrb*:TetA AmCyan^mps7^ ^13^, *TetRE*:memKillerRed^mps5^ ^13^, *pdgfrb*:CreERT2^mps6^ ^13^, *elavl3*:GFP-F^mps10^ ^11^, *elavl3*:rasmKate2^mps1^ ^13^, *elavl3*:GFP^knu3^ ^122^, *mfap4*:mCherry-F^ump6^ ^123^ (abbreviated as *mfap4*:mCherry), *mpx*:GFP^i114,73^, *mpeg1.1*:GFP-CAAX^sh425^ ^78^ (abbreviated as *mpeg1*:GFP), *14xUAS*:GCaMP6s^mpn101^ ^86^, *hsp70l*:loxp-luc2-myc-Stop-loxP-nYpet-p2a-IκBSR^ulm15^ ^124^ (abbreviated as *hs*:^loxP^Luc^loxP^IκBSR), *8xHs.NFκB*:GFP,Luciferase^hdb5^ ^125^ (abbreviated as *8xHs.NFκB*:GFP). Combinations of different transgenic zebrafish lines used in this study were abbreviated as follows: *pdgfrb*:memKillerRed (*pdgfrb*:TetA AmCyan;*TetRE*:memKillerRed), *pdgfrb*:GFP (BAC(*pdgfrb*:Gal4ff);*UAS*:EGFP), *pdgfrb*:NTR-mCherry (BAC(*pdgfrb*:Gal4ff);*UAS-E1b*:Eco.NfsB-mCherry), *pdgfrb*:GCaMP6s (BAC(*pdgfrb*:Gal4ff);*14xUAS*:GCaMP6s), *pdgfrb*:Cre^ERT2^;*hs*:^loxP^Luc^loxP^IκBSR (*pdgfrb*:CreERT2;*hsp70l*:loxp-luc2-myc-Stop-loxP-nYpet-p2a-IκBSR).

Except for scRNA-seq sample collection, AFM measurements, and functional recovery tests, embryos were treated with 0.00375% 1-phenyl-2-thiourea (PTU, Sigma-Aldrich Cat#P7629), beginning at 24 hpf, to prevent pigmentation.

For all experiments, zebrafish larvae were kept in E3 medium without methylene blue supplement ^126^.

### Zebrafish spinal cord lesions and functional recovery tests

A detailed protocol for inducing complete spinal cord lesions in zebrafish larvae has been previously described ^127^. Briefly, zebrafish larvae (72 hpf) were anesthetized in E3 medium containing 0.02% MS-222 (PharmaQ Cat#TricainePharmaQ). A 30 G x ½″ hypodermic needle was used to transect the spinal cord by either incision or perforation at the level of the urogenital pore. After surgery, larvae were returned to E3 medium for recovery and kept at 28.5 °C. Larvae that had undergone extensive damage to the notochord were excluded from further analysis. For lesions, the experimenter was blinded to the experimental treatment except for thapsigargin and BAPTA treatments, where blinding was not possible. Larvae used for lesions were randomly taken from Petri dishes containing up to 50 animals; however, no formal randomization method was used.

Analysis of behavioral recovery after spinal cord transection in larval zebrafish was performed as previously described ^25^, using Etho-Vision XT software (Noldus, version 16.0.1536). Behavioral data are shown as the distance traveled within 10 s after touch, averaged for triplicate measures per larva.

### Image Acquisition and processing

Images were acquired using the systems described in each subsection. Images and time-lapse videos were processed using ImageJ (http://rsb.info.nih.gov/ij/; version 2.3.0/1.54), Adobe Photoshop CC (version 25.5.0), Adobe Premiere Pro (version 25.0.0), and Zeiss ZEN blue software (version 3.1 blue edition). Figures were assembled using Adobe Photoshop CC.

### Whole-mount in situ hybridization (ISH)

A detailed protocol for ISH on whole-mount zebrafish larvae with digoxygenin (DIG)-labeled antisense probes has been previously described in detail ^127^. In brief, terminally anesthetized larvae were fixed in 4% formaldehyde (Thermo Fisher Scientific Cat#28908) in PBS (FA) and treated with proteinase K (Invitrogen Cat#25530-049) followed by re-fixation for 15 min in 4% FA. DIG-labeled antisense probes were hybridized overnight at 65 °C. This protocol allows efficient probe penetration in whole-mount preparations of 5 dpf larvae ^14^. Color reaction was performed after incubation with anti-DIG antibody conjugated to alkaline phosphatase (Sigma-Aldrich Cat#11093274910; RRID:AB_2313640, 1:3000) using NBT/BCIP substrate (Roche Cat#11697471001) or SIGMAFAST Fast Red (Sigma-Aldrich Cat#F4648). Samples were mounted in 75% glycerol in PBS and imaged in multi-focus mode using a Leica M205 FCA stereo-microscope equipped with a Leica DMC6200 Pixel Shift color camera, or using a Plan-Apochromat 10×/0.45 M27 objective or Plan-Apochromat 20×/0.8 objective on a Zeiss LSM 900 confocal microscope. Information on ISH probes, including primer sequences used for molecular cloning, is provided in Supplementary Data (Supplementary File 6).

### Combined hybridization chain reaction ISH and immunohistochemistry

Hybridization chain reaction (HCR) RNA fluorescence ISH (Molecular Instruments) was performed according to the manufacturer’s instructions. Information on HCR ISH probes is provided in Supplementary File 6. To enhance the GFP signal in *pdgfrb*:GFP and *elavl3*:GFP transgenic animals, we used a chicken anti-GFP antibody (Abcam Cat#ab13790; RRID:AB_300798) at a dilution of 1:1000. Samples were mounted in 75% glycerol in PBS and imaged using a Plan-Apochromat 20×/0.8 objective or C-Apochromat 40×/1.2W Korr UV-VIS-IR in tiling mode on a Zeiss LSM 980 confocal microscope.

### Immunohistochemistry (IHC)

Anti-Mpx IHC on whole-mount zebrafish larvae was carried out as previously described ^127^. Briefly, terminally anesthetized larvae were fixed in 4% FA for 1 h at room temperature. After removing the head and tail with micro scissors, larvae were permeabilized by subsequent incubation in acetone and proteinase K. Samples were re-fixed in 4% FA, blocked in PBS containing 1% Triton X-100 (PBTx; Sigma Aldrich Cat#T8787) and 4% bovine serum albumin (BSA), and incubated over three nights with rabbit polyclonal anti-Mpx antibody (GeneTex Cat#GTX128379; RRID:AB_2885768) at a dilution of 1:300.

For anti-Mfap4 IHC, terminally anesthetized larvae were fixed in 4% FA supplemented with 1% DMSO for 2 h at room temperature, followed by three washes in PBS for 5 min each. After removing the head and tail with micro scissors, larvae were incubated in 2 mg/ml Collagenase P (Roche Cat#11213857001) in PBS for 35 min. Samples were washed three times in 0.2% PBTx for 5 min each, incubated in 50 mM glycine in PBS (Sigma Aldrich Cat#50046) for 10 min, and washed again with 0.2% PBTx. Blocking was done with 1% DMSO, 2% BSA, and 0.7% PBTx in PBS for 2 h at room temperature before incubation overnight with rabbit polyclonal anti-Mfap4 antibody (GeneTex Cat#GTX132692; RRID:AB_2886714) at a dilution of 1:100 at 4°C. The next day, samples were washed 8× for 10 min in 0.2% PBTx before incubation with secondary antibody.

Secondary fluorophore-conjugated antibodies were sourced from Invitrogen and used at a dilution of 1:300 and incubated overnight at 4°C. Samples were mounted in 75% glycerol in PBS and imaged using a Plan-Apochromat 20×/0.8 objective on a Zeiss LSM 980 confocal microscope.

### Combined TUNEL/anti-GFP labeling

Combined TUNEL and anti-GFP labeling was performed as previously described ^11^. Briefly, terminally anesthetized larvae were fixed in 4% FA overnight. Larvae were permeabilized by subsequent incubation in acetone and proteinase K as described in detail elsewhere ^127^. Samples were re-fixed in 4% FA and Click-iT TUNEL Alexa Fluor 647 Imaging Assay (Thermo Fisher Cat#C10247) was performed according to the manufacturer’s protocol to label apoptotic cells. Following labeling, samples were blocked in 3% BSA in PBTx (1% Triton-X in PBS) and incubated over three nights with chicken anti-GFP antibody (Abcam Cat#ab13790; RRID:AB_300798, 1:500). After several washes in PBTx, samples were incubated over two to three nights with the secondary antibody of interest (Invitrogen, 1:300 diution). Following secondary labeling, samples were washed in PBTx and mounted in 75% glycerol in PBS. Imaging was done using a Plan-Apochromat 10×/0.45 M27 objective on a Zeiss LSM 980 confocal microscope.

### Live imaging of larval zebrafish

For live confocal imaging, zebrafish larvae were anesthetized in E3 medium containing 0.02% MS-222 and mounted in the appropriate orientation in 1% low-melting-point agarose (Ultra-PureTM Low Melting Point, Invitrogen Cat#16520) between two microscope cover glasses (Menzel, thickness I) or in a glass-bottom petri dish (ibidi Cat#81158). During imaging, larvae were covered with 0.01% MS-222-containing E3 medium to keep preparations from drying out. Imaging was done using a Plan-Apochromat 10×/0.45 M27 objective, Plan-Apochromat 20×/0.8 objective, and C-Apochromat 40×/1.2W Korr UV-VIS-IR objective on a Zeiss LSM 980 confocal microscope, or using a Leica M205 FCA stereomicroscope equipped with a Leica DMC6200 Pixel Shift color camera.

### Optoablation

For optoablation, *pdgfrb*:memKillerRed transgenic zebrafish larvae were anesthetized in E3 medium containing 0.02% MS-222 and mounted in a lateral position in 1% low-melting-point agarose between two microscope cover glasses, filled with 0.01% MS-222-containing E3 medium. Larvae were irradiated with intense green light from a mercury lamp (HXP200C, Zeiss filter set 63 HE mRFP with EX BP 572/25 nm, 144 mW; HXP120V, Zeiss filter set 43 Cy3 with EX BP 545/25 nm, 160 mW) for 30 min each, on a Zeiss LSM 980 (Plan-Apochromat 20×/0.8 objective) or Zeiss Axio Zoom.V16 microscope (PlanNeoFluar 2.3× objective and 258× zoom). Irradiation by both set-ups resulted in comparable bleaching of most of the memKillerRed fluorescence signal in approximately 1 mm trunk tissue surrounding the urogenital pore.

### Drug treatments and heat shocks

Drug treatments and heat shocks were performed according to the schematic timelines shown with each experiment. Larvae were incubated in E3 medium containing the drug. To deplete intracellular Ca^2+^, we used the cell-permeant chelator BAPTA-AM (Thermo-Fisher Cat#B1205) dissolved in DMSO at a final concentration of 50 µM, previously described for use in zebrafish ^87^. To deplete Ca^2+^ in the endoplasmic reticulum store, we used thapsigargin (Sigma-Aldrich Cat#586005) dissolved in DMSO at a final concentration of 1 µM, as previously described in zebrafish larvae ^88^. Larvae that developed severe heart edema or other malformations at 24 hpl, were excluded from analysis. The NFκB inhibitor JSH-23 (Sigma-Aldrich Cat#J4455) was dissolved in DMSO and used at a final concentration of 5 µM. Doxycycline (DOX; Sigma-Aldrich Cat#M3761) treatments to induce memKillerRed expression in *pdgfrb*:memKillerRed transgenic animals were performed as previously described in detail elsewhere ^13^. Briefly, DOX was dissolved in reverse osmosis H_2_O and used at a final concentration of 25 µg/ml. DOX was added to the E3 medium at 24 hpf. 4–hydroxytamoxifen (4OHT; Sigma-Aldrich Cat#H7904) treatments for Cre^ERT^^2^-mediated recombination were performed as previously described in detail elsewhere ^13^. Briefly, 4OHT was dissolved in 100% ethanol and used at a final concentration of 10 mM. Controls refer to 4OHT-treated clutch mates of either *pdgfrb*:Cre^ERT2^ or *hs*:^loxP^Luc^loxP^IκBSR transgenic alleles. For heat shocks, larvae were kept in 50-ml conical centrifuge tubes filled with E3 medium, which floated in a programmable thermostat-controlled water bath (Lauda, Germany). Heat shocks were performed for 1 h at 38° C after which larvae were returned to 28.5°C. For targeted cell ablation of *pdgfrb^+^* cells, the pro-drug Metronidazole (MTZ; Sigma-Aldrich Cat#M3761) was dissolved in DMSO and used at a final concentration of 2 mM. MTZ was added to the E3 medium, beginning at 48 hpf. Larvae that developed severe heart or brain edema were excluded from analysis. Lipopolysaccharides from *Escherichia coli* (LPS; Sigma-Aldrich Cat#L2880) was dissolved in PBS and used at a final concentration of 50 µg/ml. The Cxcr1/2 inhibitor SB225002 (Sigma-Aldrich Cat#559405) was dissolved in DMSO and used at a final concentration of 2.5 µM.

### Morpholino treatments

Custom-designed Morpholinos (Gene Tools) were dissolved in nuclease-free H_2_O at a concentration of 1 mM. Amounts injected into the zygote and sequences of the splice blocking (splice MO) and translation blocking MO (ATG MO) used are as follows: *cxcl8b.1* ATG MO: 5ʹ-AGGCTGAAACGCTCAACTTCATCAT-3′ (8 ng), *tgfb1a* ATG MO: 5′-TCAGCACCAAGCAAACCAACCTCAT-3′ (4 ng), and *cxcl8b.1* splice MO: 5′-GTTAGTATCTGCTTACCCTCATTGG -3′ (4 ng). *tgfb1a* ATG MO and *cxcl8b.1* splice MO have previously been established in zebrafish ^91–93^. As control, a standard MO from Gene Tools targeting a mutated splice site of human β-Globin gene was used: 5′-CCTCTTACCTCAGTTACAATTTATA-3′ (4 ng).

### CRISPR/Cas9 manipulation

CRISPR/Cas9-mediated mutagenesis was achieved following a previously published protocol with minor modifications ^128^. Three gRNAs targeting exon 1 or 2 of the *tgfb1a* gene (ENSDARG00000041502) were designed using the CHOPCHOP webtool (https://chopchop.cbu.uib.no/) ^129^: gRNA#1 target site: 5ʹ-ATGGCTAAAGAGCCTGAATCCGG-3ʹ (exon 1), gRNA#2 target site: 5ʹ-CTGGAACTGTATCGCGGAGTGGG-3ʹ (exon 2), gRNA#3 target site: 5ʹ-TCGGATCAAGAATCCCACCATGG-3ʹ (exon 2). Custom crRNAs (Alt-R™ CRISPR-Cas9 crRNA, IDT) and tracrRNA (Alt-R® CRISPR-Cas9 tracrRNA, IDT Cat#1072532) were resuspended in Duplex buffer (IDT Cat#11-01-03-01) to a concentration of 200 µM and annealed at 95°C for 5 min to obtain gRNAs. gRNA#1–3 were pooled and incubated at 37°C for 5 min with recombinant Alt-RT^TM^ S.P.Cas9 Nuclease (IDT Cat#107807) to assemble the gRNA/Cas9 ribonucleoprotein complex (RNP). A total of 75 pg of each gRNA and 1 ng Cas9 protein were microinjected into the zygote. Negative control crRNA (Alt-R® CRISPR-Cas9 Negative Control crRNA #1–3, IDT Cat#1072544, Cat#1072545, Cat#1072546) was used as control. Efficient mutagenesis of the targeted loci was confirmed with melting curve analysis (SYBR® Green, Thermo Fisher Cat#A25918). The following oligos were used to amplify a fragment of genomic DNA containing the respective target sites: gRNA#1, 5ʹ-AGCATTATGAGGTTGGTTTGCT-3ʹ and 5ʹ-TTCATTTCTTCGCTCAGTTCAA-3ʹ; gRNA#2/3, 5ʹ-TCAGATGTTTTTCAACGTGTCC-3ʹ and 5ʹ-CTGTTTCACGTCAAATGAGAGC-3’.

### Tissue dissociation and cell sorting

For single-cell transcriptomics (scRNA-seq) covering the entire regeneration time course (scRNA-seq dataset #1), trunk tissue of *pdgfrb*:GFP transgenic zebrafish larvae was isolated at 6, 12, 24, and 48 hpl as well as from uninjured controls. All samples were collected in duplicates and age-matched (4 dpf) except for the 48 hpl samples, which were from 5 dpf animals. For scRNA-seq of the lesion site from control and fibroblast-depleted animals (scRNA-seq dataset #2), trunk tissue of *pdgfrb*:NTR-mCherry transgenic larvae treated with either MTZ or DMSO control was isolated. Samples were collected in duplicates. Zebrafish larvae were terminally anesthetized in E3 medium containing 0.02% MS-222 and transferred to ice-cold 1× Hank’s Balanced Salt Solution in H_2_O (HBSS [10×]; Thermo Fisher Cat#14065056). Micro scissors were used to isolate trunk tissue spanning approximately four somites in length and containing the lesion site or corresponding unlesioned tissue. Trunk tissue was collected in ice-cold 1× HBSS. Per sample, excised tissue from 125 larvae was pooled and enzymatically dissociated at 37°C using 1 ml 0.25% Trypsin-EDTA (Thermo Fisher Cat#25200072) followed by mechanical dissociation using a fire-polished glass Pasteur pipette. Enzymatic digestion was terminated by adding 10 ml ice-cold 1× HBSS. Cell suspension was filtered through a 20-µm cell strainer (pluriSelect Cat#43-50020-03). Following centrifugation for 10 min at 300 g and 4°C, cells were resuspended in 500 µl 1× HBSS containing 50 µM calcein (Thermo Fisher Cat#C1429). Viable cells were isolated by FACS using 405 nm excitation and a BV421 filter at a MoFlo Astrios EQ (Beckman Coulter) or BD FACSAria II (BD Biosciences) device. Up to 100,000 calcein^+^ cells were sorted into 1× HBSS to obtain a density of ∼700 cells/µl.

### Single-cell library preparation and RNA sequencing

Single cells were captured using a Chromium Single Cell 3’ kit (v3.1 chemistry, 10x Genomics Cat#PN-1000128) according to manufacturer’s instructions, aiming for 10,000 cells per library. Complementary DNA was amplified for 11 cycles and index primer addition PCR used 12–14 cycles, depending on the amount of cDNA. Sequencing of the libraries was performed using the Illumina NovaSeq 6000 S2 flow cell, running 500 million reads/fragments in paired-end (asynchronous) mode per sample (i.e., ∼50,000 reads/cell).

### Quality control and clustering of scRNA-seq data

To prepare a custom zebrafish transcriptome, FASTA sequences coding for GFP and Gal4ff were appended to the *Danio rerio* genome assembly GRCz11, which was indexed with v4.3.2 zebrafish transcript annotations using CellRanger v7.1 software ^130,131^. CellRanger output matrices were analysed using the Scanpy package v1.9.1 ^132^. Cells possessing fewer than 200 unique molecular identifiers (UMIs), more than 5% mitochondrial genes, or over 5,000 counts were filtered out in quality control steps. We also excluded genes which were found to be expressed in fewer than 3 cells. Doublets were predicted with the Scrublet package using the raw (unnormalized) count matrix and removed from the data ^133^. After merging of all the samples, normalization and log-transformation, data was corrected for batch-to-batch variations using the BBKNN algorithm ^134^. Graph-based clustering was done using the parameters indicated in Supplementary File 7. Differential gene expression was calculated using Wilcoxon rank-sum test. Clusters were annotated according to known cell type markers in zebrafish or orthologous markers in mammals among the top differentially expressed genes per cluster (Supplementary File 1).

### Downstream analysis of scRNA-seq data

Mean cell numbers of fibroblast-like subclusters (scRNA-seq dataset #1) were normalized to the total number of fibroblasts in the respective condition. Clusters fulfilling all three of the following criteria were considered injury-specific fibroblast states: ≤ 1% unlesioned cells, foldchange over unlesioned ≥ 2 for two out of 6, 12, and 24 hpl conditions, ≤ 2% 48 hpl cells.

RNA velocity was estimated using velocyto package v0.17.15 to annotate spliced/unspliced reads and scVelo v0.2.4 (dynamical model) ^135,136^.

Gene Ontology (GO) Enrichment Analysis for biological process, molecular function and cellular component was performed using the PANTHER based http://geneontology.org website ^137–139^.

To predict cell cycle phases, zebrafish orthologues of human cell cycle genes were compiled using the HGNC Comparison of Orthology Predictions (HCOP) tool (see Supplementary File 4) ^140,141^. Cell cycle phases were calculated by computing S phase and G2/M phase scores in Scanpy. Mean numbers of cells in G1, S or G2/M phase were normalized to the total number of cells from the respective condition (i.e. time point after SCI).

Human reference data for pathway activity inference was obtained from PROGENy, zebrafish orthologues for human target genes were obtained from g:Profiler, and mean pathway activity per cluster was calculated using decoupler-py v1.5.0 (see Supplementary File 8) ^142–144^.

The interactome of *cthrc1a^+^* fibroblasts and neutrophils/Mϕ was predicted using the DanioTalk R package ^145^, which is combining physical interaction data of the zebrafish proteome with existing human ligand-receptor pair databases. Minimum gene expression cutoff was set to 0.01, fold-change data was included to rank interactions.

Mean cell numbers of neutrophil and Mϕ subclusters in scRNA-seq dataset #1 were normalized to the total number of hematopoietic cells in the respective condition to illustrate their relative abundance in the lesion core over time. Mean cell numbers of neutrophil and Mϕ subclusters in scRNA-seq dataset #2 were normalized to the total number of neutrophils or Mϕ in the respective condition to illustrate the shift in the relative abundancies of the different neutrophil and Mϕ subpopulations.

### Cross-polarized optical coherence tomography (CP-OCT)

For *in vivo* CP-OCT ^95^, zebrafish larvae were anesthetized in E3 medium containing 0.02% MS-222 and mounted in a lateral position in 1% low-melting-point agarose between two microscope cover glasses. During imaging, larvae were covered with 0.01% MS-222-containing E3 medium to keep preparations from drying out. Cross-polarized images of the spinal lesion site were acquired using a custom-built CP-OCT system ^146^. Light from a broadband supercontinuum laser (YSL Photonics Cat#SC-OEM) was filtered to acquire a spectrum centered at 885 nm with a full width at half maximum of 80 nm. The laser was operated at 200 MHz. The filtered spectrum from the laser was coupled to a single-mode optical fiber and collimated using a collimator (Thorlabs Cat#F230APC-850). The light was further split using a 90/10 beam splitter (Thorlabs Cat#BS025) into a reference beam and a sample beam. The sample was illuminated with 15 mW of optical power. A combination of a quarter-wave plate (Thorlabs Cat#SAQWP05M-1700) and a lens (Thorlabs Cat#AC254-030-AB) was inserted into the reference and the sample arm to control the polarization of the light. A galvano mirror (Thorlabs Cat#GVS012) was used to scan the laser beam over the sample. The reflected reference and the sample signals were acquired using a custom-designed spectrometer consisting of a reflective collimator (Thorlabs Cat#RC08APC), a holographic grating (Wasatch Photonics Cat#1200 l/mm@840nm), a lens (Thorlabs Cat#AC-254-080-B), and a line scan camera (Basler Cat#2048 pixels, ral2048-48gm) operating at 25 kHz line scan rate. The spectrometer signal was processed using LabVIEW-based custom-designed software (National Instruments). The acquired interference spectrum from the camera was spectrally recalibrated from wavelength space to wavenumber space ^147^ and a fast Fourier transform was performed to obtain an axial profile of the sample.

### Combined confocal fluorescence and Brillouin microscopy (BM)

For *in vivo* BM, zebrafish larvae were anesthetized in E3 medium containing 0.02% MS-222 and mounted in a lateral position in 1% low-melting-point agarose on a 35 mm glass-bottom dish (ibidi Cat#81158). During imaging, larvae were covered with 0.01% MS-222-containing E3 medium to keep preparations from drying out. Brillouin frequency shift images were acquired by BM, employing a confocal configuration and a Brillouin spectrometer consisting of a two-stage virtually imaged phase array (VIPA) etalon, as previously described in detail elsewhere ^148^. Briefly, the sample was illuminated by a frequency-modulated diode laser beam (*λ* = 780.24 nm, DLC TA PRO 780, Toptica), which was stabilized to the D_2_ transition of rubidium ^85^Rb. A Fabry–Perot interferometer in two-pass configuration and a monochromatic grating (Toptica) were employed to further suppress the contribution of amplified spontaneous emission to the laser spectrum. The laser light was coupled into a single-mode fiber and guided into the backside port of a commercial inverted microscope stand (Axio Observer 7, Zeiss). An objective lens (20×, NA = 0.5, EC Plan-Neofluar, Zeiss) illuminated the sample on a motorized stage with an optical focus. The laser power at the sample plane was set at 15 mW. The backscattered light from the sample was collected by the same objective lens, coupled into the second single-mode fiber to achieve confocality, and delivered to a Brillouin spectrometer. In the Brillouin spectrometer, the backscattered light was collimated and passed through a molecular absorption cell filled with rubidium ^85^Rb (Precision Glassblowing Cat#TG-ABRB-I85-Q), in which the intensity of the Rayleigh scattered and reflected light was significantly suppressed. After passing through the molecular absorption cell, the beam was guided to two VIPA etalons (Light Machinery Cat#OP-6721-6743-4) with the free spectral range of 15.2 GHz, which convert the frequency shift of the light into the angular dispersion in the Brillouin spectrum. The Brillouin spectrum was acquired by a sCMOS camera (Teledyne Cat#Prime BSI), with the exposure time of 0.5 s per measurement point. The two-dimensional Brillouin frequency map of the injured region was measured by scanning the motorized stage on the microscope stand, with the translational step size of 0.5 µm. The Brillouin microscope was controlled with custom acquisition software written in C++ (https://github.com/BrillouinMicroscopy/BrillouinAcquisition; version 0.3.4). Confocal fluorescence imaging was performed in the same region of interest as Brillouin measurement using a re-scan confocal microscopy (RCM) module (RCM2, Confocal.nl) that was attached to one side port of the microscope stand. The RCM module consists of a sCMOS camera (Prime BSI Express, Teledyne) and a multi-line laser unit (Skyra, Cobolt) as an excitation illumination source for four laser lines (*λ* = 405, 488, 562, and 637 nm). A pinhole of the diameter of 50 µm and the re-scanning imaging principle ^149^ provides rapid confocal fluorescence imaging with a lateral resolution of 120 nm. By focusing a confocal fluorescence image in the center of the spinal cord of *elavl3*:GFP-F transgenic zebrafish larvae, the axial plane of Brillouin imaging was determined.

### Optical diffraction tomography (ODT)

For *in vivo* ODT, zebrafish larvae were anesthetized in E3 medium containing 0.02% MS-222 and mounted in a lateral position in 1% low-melting-point agarose on a 35 mm glass-bottom dish (ibidi Cat#81158). During imaging, larvae were covered with E3 medium containing 0.01% MS-222 and 20% of refractive index (RI)-matching agent iodixanol (OptiPrep^TM^; Sigma-Aldrich Cat#D1556) ^150^. Iodixanol was used to reduce the RI difference between the zebrafish larvae and the surrounding medium. The final RI of the medium was 1.351, which was determined by an Abbe refractometer (Kern & Sohn GmbH Cat#ORT1RS). The RI distribution of the zebrafish spinal lesion site was measured by ODT, employing Mach-Zehnder interferometry to measure multiple complex optical fields from various incident angles, as previously described ^151^. A solid-state laser beam (*λ* = 532 nm, 50 mW, CNI Optoelectronics Technology Co.) was split into two paths using a beam splitter. One beam was used as a reference beam and the other beam illuminated the sample on the stage of an inverted microscope (Axio Observer 7, Zeiss) through a tube lens (*f* = 175 mm) and a water-dipping objective lens (40×, NA = 1.0, Zeiss). A high numerical aperture objective lens (40×, water immersion, NA = 1.2, Zeiss) collected the beam diffracted by the sample. To reconstruct a 3D RI tomogram of the sample, it was illuminated from 150 different incident angles scanned by a dual-axis galvano mirror (Thorlabs Cat#GVS212/M) located at the conjugate plane of the sample. The diffracted beam interfered with the reference beam at an image plane and generated a spatially modulated hologram, which was recorded with a CMOS camera (XIMEA Cat#MQ042MG-CM-TG). The field-of-view of the camera covers 205.0 μm × 205.0 μm. The complex optical fields of light scattered by the samples were retrieved from the recorded holograms by applying a Fourier transform-based field retrieval algorithm ^152^. The 3D RI distribution of the samples was reconstructed from the retrieved complex optical fields via the Fourier diffraction theorem, employing the first-order Rytov approximation ^153,154^. A more detailed description of tomogram reconstruction can be found elsewhere ^155^. The MATLAB script for ODT reconstruction can be found at https://github.com/OpticalDiffractionTomography/ODT_Reconstruction (version 1.0.0).

### Atomic force microscopy (AFM)-based indentation measurements

*In vivo* AFM measurements of the zebrafish lesion core were performed using silicon cantilevers (Arrow-TL1, NanoWorld) with a custom-attached polystyrene bead as probe (∼37 µm diameter, microParticles GmbH) mounted on a JPK Nanowizard Cellhesion 300 (JPK Instruments AG). Cantilever spring constants were determined via the thermal noise method and cantilevers with spring constants of 0.02–0.04 N/m selected for measurements ^156^. Terminally anesthetized larvae were mounted in an upright position in 3% low-melting-point agarose. The trunk tissue was cut transversely using a vibratome to expose the lesion core. The agarose block containing the larva was glued onto a plastic tissue culture dish. During measurements, larvae were covered with 0.01% MS-222 in PBS to keep preparations from drying out. The ROI for indentation was identified by fluorescence signal from the rostral spinal cord stumps in *elavl3*:GFP-F transgenics using a custom-built upright setup with an Axio Zoom.V16 (Zeiss) fluorescence microscope with a pco.edge 4.2 bi sCMOS camera. Force–distance curves were taken with a maximum force of 5 nN at an approach speed of 7 µm/s.

### Quantifications and statistics

For quantification of axonal bridge thickness, transgenic live animals (*elavl3*:GFP-F or *elavl3*:rasmKate2) were imaged using a Plan-Apochromat 10×/0.45 M27 objective on a Zeiss LSM 980 confocal microscope in lateral view. The length of a vertical line that covers the width of the axonal bridge at the center of the lesion site, was then determined in maximum intensity projections using ImageJ software (https://imagej.nih.gov/ij/index.html). Except for Supplementary Fig. 5i, measurements were performed in samples of at least three independent experiments and spinal cord lesions for at least one out of the experimental replicates was induced by a second operator.

For quantification of neutrophils and Mϕ in the lesion core, transgenic live animals (*mpx*:GFP, *mfap4*:mCherry, *mpeg1*:GFP) or whole-mount anti-Mpx or anti-Mfap4 IF samples were imaged using a Plan-Apochromat 20×/0.8 objective on a Zeiss LSM 980 confocal microscope in lateral view. Quantification of cells was performed with ImageJ software in a pre-set ROI of constant size (150 µm × 50 µm) on individual optical sections of a confocal stack. The ROI was ventrally limited by the notochord.

For analysis of neutrophil and Mϕ recruitment to the lesion core in time-lapse videos, *mpx*:GFP;*mfap4*:mCherry double transgenic animals were imaged using a Plan-Apochromat 10×/0.45 M27 objective on a Zeiss LSM 980 confocal microscope in lateral view. Quantification of cells was performed with ImageJ software in a pre-set ROI of constant size (*r* = 50 µm) on maximum intensity projections of a confocal stack. The ROI was placed in the center of the lesion. We did not consider cells that were present in or immediately surrounding the ROI before the SCI.

Quantitative analysis of fluorescent ISH signals in the lesion site was performed on captured images of whole-mount samples following previously published protocols ^14,15^. Briefly, the number of pixels above an intensity threshold (thresholded pixel area) in a ROI was determined on maximum intensity projections using ImageJ software. The intensity threshold was set individually for each experimental dataset, using untreated lesioned control samples as a reference. The ROI was ventrally limited by the midline.

Extraction and quantification of Brillouin frequency shifts were performed in a pre-set ROI, using custom software written in Python (https://github.com/GuckLab/impose, version 0.4.10; https://github.com/BrillouinMicroscopy/BMicro, version 0.8.2). The ROI was 40 µm × 20 µm, which corresponds to the determined average axonal bridge thickness and the average distance between the spinal cord stumps following a dorsal incision lesion at 1 dpl. The ROI was placed in the center of the lesion site and ventrally limited 5 µm dorsal to the notochord. Brillouin frequency shifts were measured in samples of six independent experiments. The measured Brillouin frequency shift *v*_B_ can be expressed in terms of the longitudinal modulus *M*′, refractive index (RI) *n*, and density *ρ* of the specimen, as well as the incident wavelength *λ*_0_ and scattering angle 𝜗 given by the setup: 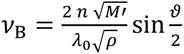. All measurements were performed in the backscattering configuration with 𝜗 = 180° and, accordingly 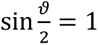. The longitudinal compressibility *κ_L_* may be expressed as the inverse of the longitudinal modulus 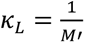. For the sake of conciseness *κ_L_* will be referred to as ‘compressibility’, despite the fact that this term is usually defined as the inverse of the bulk modulus *K*′. Bulk modulus *K*′, shear modulus *G*′, and longitudinal modulus *M’* are related by the following equation in the case of an isotropic sample: 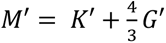.

To quantify the co-polarization ratio (ratio of preserved polarization to total reflectivity) in the lesion site, two images of the tissue were acquired: one co-polarized and one cross-polarized image. For the acquisition of the co-polarized image, the angle of the quarter-wave plate (QWP) axis in the reference path was set in such a way that the light in the sample and the reference arm had the same polarization. To acquire the cross-polarized image, the QWP in the reference beam path was rotated by 45°, resulting in orthogonal polarization between reference and sample arm. Sample reflectivity *R*(*z*) and co-polarization ratio *δ*(*z*) of the sample was calculated using the amplitude of the co-(*A_co_*) and cross-(*A_cross_*) polarized images using the following equations: 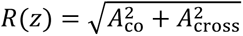 and 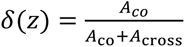. Using ImageJ software, the co-polarization ratio was determined in a pre-set ROI of constant size (80 µm × 40 µm) for the averaged intensity projection of the complete image stack. The ROI was placed in the center of the lesion site and ventrally limited by the notochord. The results were validated by a second independent observer.

Extraction and quantification of the RI and mass density distribution from reconstructed tomograms were performed using custom-written MATLAB (MathWorks, version R2018b) scripts. The mass density of the samples was calculated directly from the reconstructed RI tomograms, since the RI of the samples, *n*(*x*, *y*, *z*), is linearly proportional to the mass density of the material, *ρ*(*x*, *y*, *z*), as *n*(*x*, *y*, *z*) = *n*_m_ + *αρ*(*x*, *y*, *z*), where *n*_m_ is the RI value of the surrounding medium and *α* is the RI increment (*dn*/*dc*), with *α* = 0.19 ml/g for proteins and nucleic acids ^157,158^. RI and mass density distribution from reconstructed tomograms were evaluated in a pre-set ROI in a sagittal slice of the RI tomogram along the *x*-*y* plane at the focused plane, using a custom-written MATLAB script. The ROI was the same as applied to Brillouin images. Note that the RI was measured in samples independent of those analyzed with BM.

AFM indentation analysis was performed using a custom-written MATLAB script (Batchforce; MathWorks, https://github.com/FranzeLab/AFM-data-analysis-and-processing). The Hertz model was applied to fit the data and extract the apparent elastic modulus *K* ^159^. Force-indentation curves that could not be fitted to the Hertz model were excluded from analysis.

For all quantifications, investigators were blinded to the experimental conditions. Unless indicated, no data were excluded from analysis. All statistical analyses were performed using Graph Pad Prism 10 (GraphPad Software Inc., version 10.3.1). All quantitative data were tested for normal distribution using Shapiro-Wilk test. Parametric and non-parametric tests were used as appropriate. Statistical tests used in each experiment are given in the figure legends. We used two-tailed Student’s *t*-test, two-tailed Mann– Whitney test, Fisher’s Exact, and Kruskal–Wallis test followed by One-way Anova with Šídák’s multiple comparisons test. Differences were considered statistically significant at *p*-values below 0.05. Unless otherwise indicated, variance of all group’s data is presented as ± standard error of the mean (SEM). The sample size (*n*) for each experimental group is given in the figure legends. Graphs were generated using GraphPad Prism 10.

## DATA AVAILABILITY

Except for the scRNA transcriptomics data, all data are available in the main manuscript, or the supplementary materials. The scRNA transcriptomics data reported in this paper can be explored under https://shiny.mpl.mpg.de/wehner_lab/spinal_cord_regeneration_atlas/ with an interactive dashboard based on the iSEE software and have been deposited at NCBI Gene Expression Omnibus (GEO; https://ncbi.nlm.nih.gov/geo/) under the following accession numbers: XX (scRNA-seq dataset #1) and XX (scRNA-seq dataset #2) ^160^. Source data is provided with this paper.

## ACKNOWLEDGEMENTS

We thank Dr. Eoghan O’Connel for advice on data analysis with Python, Raül González Garrido for support in creating the web ressource, Drs Catherina Becker and Thomas Becker for critical reading, Casandra Carrillo Mendez, Olga Stelmakh, Heidi von Berg, and Tina Locker for excellent fish care. This work was supported by the Animal Facility of the Max-Planck-Zentrum für Physik und Medizin, the Core Unit Bioinformatics of the Universitätsklinikum Erlangen, the Core Unit für Zellsortierung und Immunomonitoring of the Friedrich-Alexander-Universität Erlangen-Nürnberg, and the Sequencing facility of the Max Planck Institute for Molecular Genetics. The authors acknowledge financial support from the Max Planck Society (to J.G.), the European Research Council (Synergy Grant 101118729 UNFOLD to K.F.), the Alexander von Humboldt Foundation (Alexander von Humboldt Professorship to K.F.), the Deutsche Forschungsgemeinschaft (project number 527729149 to D.W.; project number 31834696 – SFB1292 TP19N to F.M.; project number 460333672 – CRC1540 Exploring Brain Mechanics to D.W., J.G., K.F., S.F., project number 270949263 – GRK2162 to K.F.), the Bavarian State Ministry of Sciences, Research, and the Arts (ForInter; F.2-F2412.30/1/24 to S.F.), and the Interdisciplinary Center for Clinical Research (IZKF) at the University Hospital of the FAU Erlangen-Nürnberg (P074, E32 to S.F.).

## AUTHOR CONTRIBUTIONS STATEMENT

(CRediT nomenclature)

N.J.: formal analysis, investigation, visualization, writing – original draft, writing – review & editing; T.F.: formal analysis, investigation, visualization, writing – original draft, writing – review & editing; J.K.: investigation; O.L.: formal analysis, investigation; S.V.-S.: formal analysis, investigation; A.P.: formal anaylsis, investigation; K.K.: formal analysis; investigation; M.T.: investigation; P.G.: formal analysis, data curation; K.S.: resources, supervision; F.M.: data curation, visualization; S.S.: data curation, visualization; S.F.: formal analysis; K.F.: resources, supervision, funding acquisition; formal analysis; J.G.: resources, supervision, funding acquisition; D.W.: conceptualization; project administration, formal analysis, investigation, visualization, writing – original draft, writing – review & editing, resources, supervision, funding acquisition.

## Supplementary Information

**Supplementary Figure 1.**
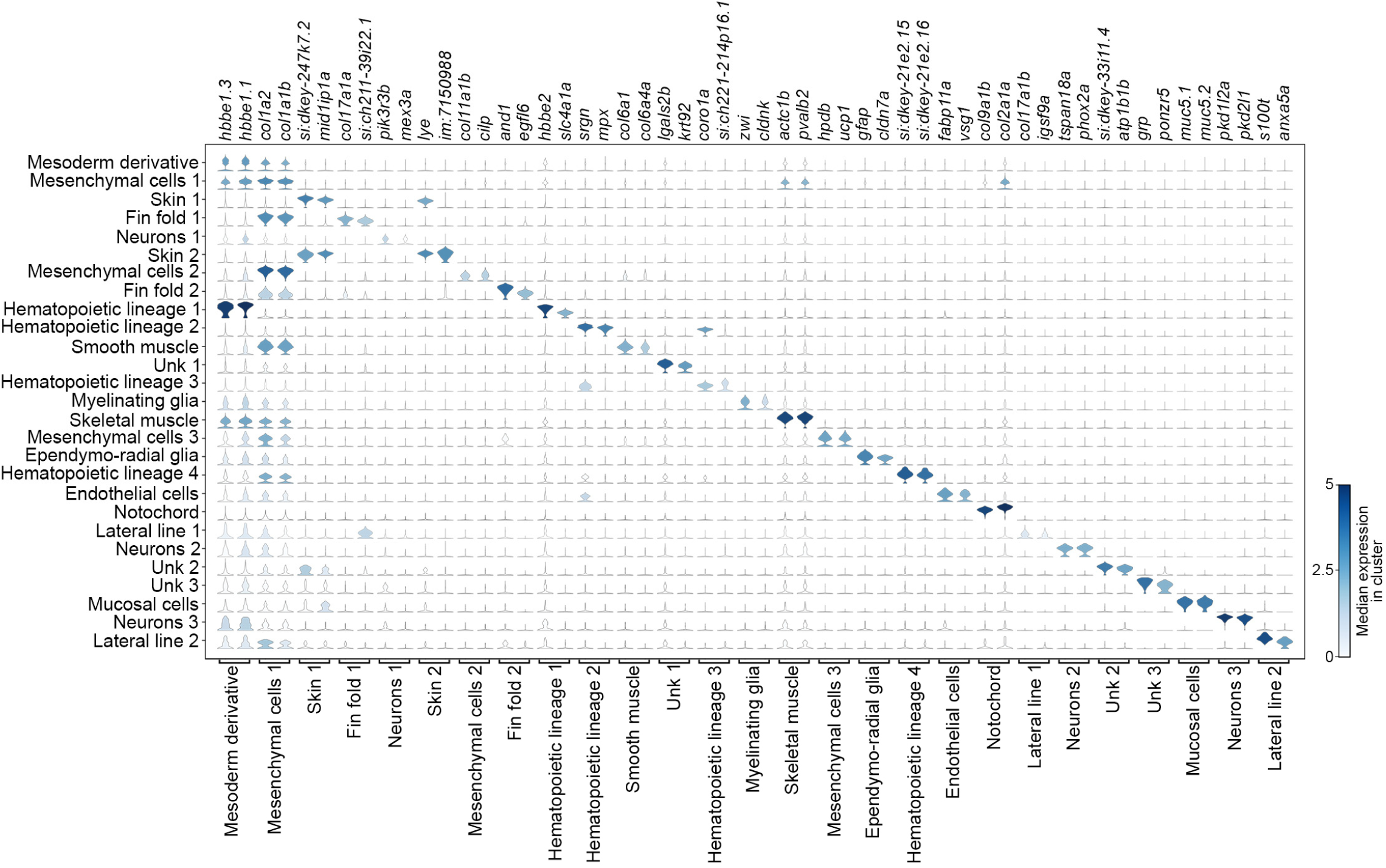
Single-cell transcriptome analysis of zebrafish spinal cord regeneration. Violin plots displaying expression of indicated genes in each major cluster presented in Fig. 1d. Abbreviation: unk, unknown.

**Supplementary Figure 2.**
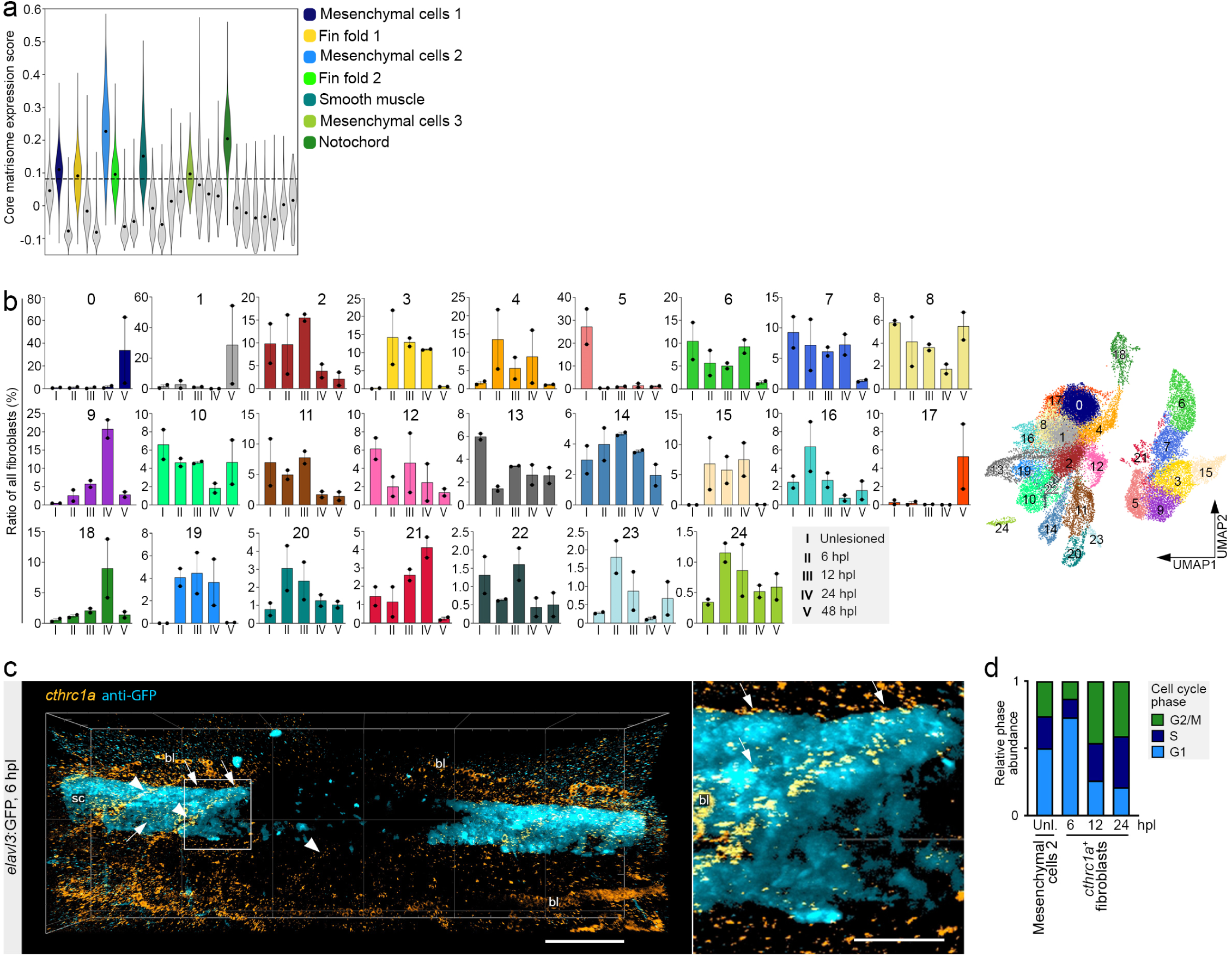
Identification of injury-specific transcriptional fibroblast states. **a)** Expression score of core matrisome genes of individual major clusters presented in Fig. 1. Data points represent means. Clusters exhibiting at least three-fold higher score than the mean score of all clusters (dashed line) were considered as fibroblast-like and are highlighted in color. **b)** Cell number dynamics over all timepoints and UMAP representation of fibroblast-like subclusters. Each data point represents one independent biological replicate, normalized to the total number of fibroblasts per sample. Data are means ± SEM. **c)** Expression of *cthrc1a* (orange) is upregulated in regions surrounding the spinal cord (cyan; arrows) and in myotendinous junctions near and distant to the wound edge (arrowheads) at 6 hpl. Image shown is a 3D reconstruction of the specimen presented in Fig. 2e. Scale bars: 100 µm and 50 µm (inset). **d)** *cthrc1a*^+^ fibroblasts exhibit enhanced proliferation at the 12 and 24 hpl timepoints. Relative abundance of cells in G1, S and G2/M cell cycle phases (determined by cell cycle gene expression) in indicated clusters and conditions is shown. **a**–**d)** Abbreviations: bl, blood; hpl, hours post-lesion; UMAP, uniform manifold approximation and projection; unl, unlesioned.

**Supplementary Figure 3.**
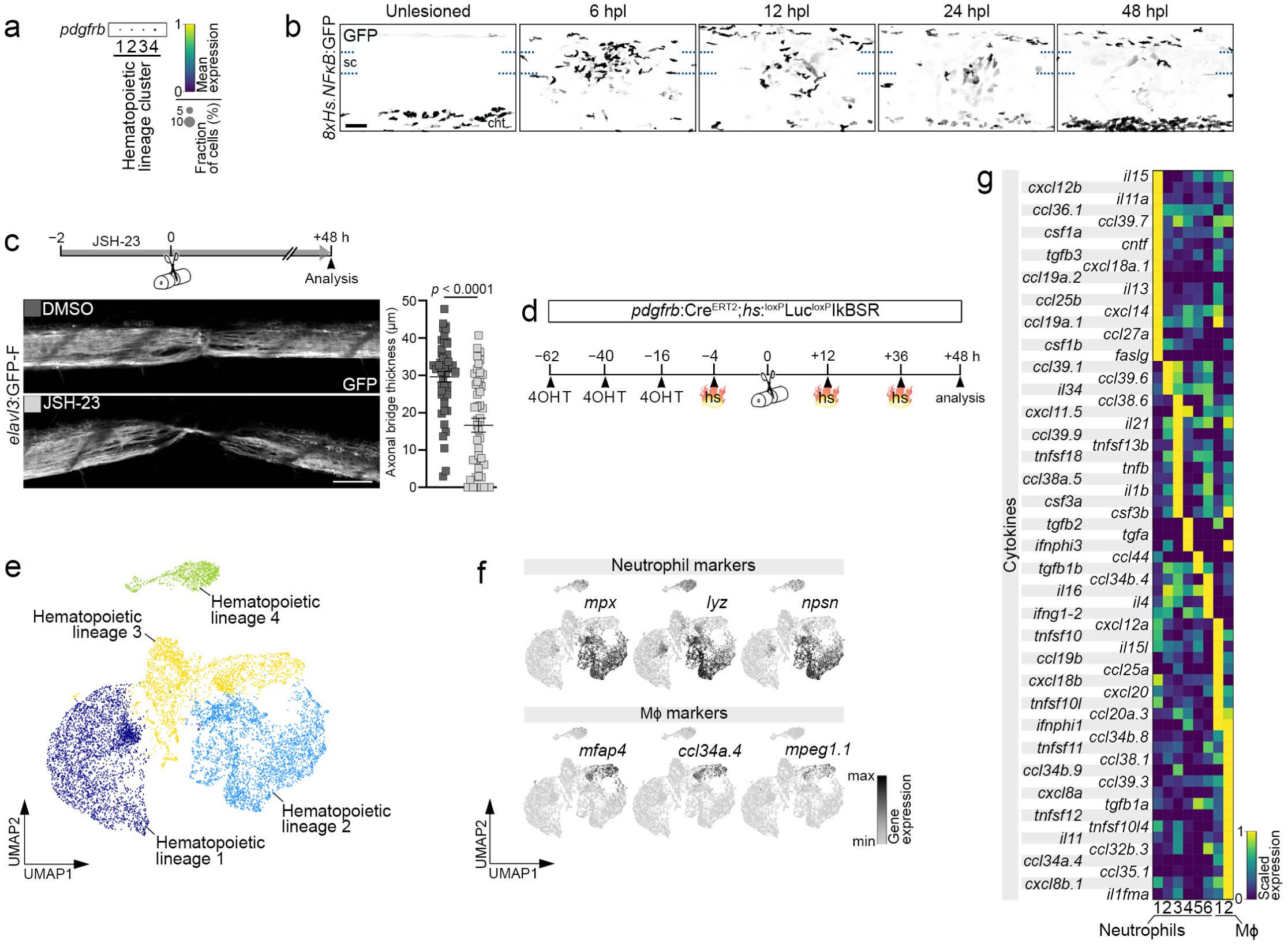
*cthrc1a*^+^ fibroblasts have an inflammatory profile and characterization of macrophage and neutrophil populations. **a)** Mean expression of *pdgfrb* in indicated hematopoietic lineage clusters. Note: *pdgfrb* transcripts are not detectable in the vast majority of hematopoietic lineage cells. **b)** Cells with high NFκB pathway reporter activity (GFP^+^, black) rapidly accumulate in the lesion site, likely representing neutrophils of caudal hematopoietic tissue (cht) origin. Images shown are maximum intensity projections of the unlesioned trunk or lesion site (lateral view; rostral is left). Dashed lines indicate location of the spinal cord. *n* ≥ 10 for each timepoint. **c)** Treatment with the NFκB inhibitor JSH-23 reduces the thickness of the axonal bridge (white) at 48 hpl. Images shown are maximum intensity projections of the lesion site (lateral view; rostral is left). Data are means ± SEM. Each data point represents one animal. Two-tailed Mann–Whitney test. *n* = 48 for each condition over three independent experiments. **d)** Detailed timeline of 4-hydroxytamoxifen (4OHT) treatment and heat shock-induced (hs) expression of IκBSR in *pdgfrb*:Cre^ERT2^;*hs*:^loxP^Luc^loxP^IκBSR transgenic animals of the experiment shown in Fig. 3e. **e)** UMAP representation of all hematopoietic lineage clusters. **f)** UMAP representation of all hematopoietic lineage cells showing scaled expression (black) of commonly accepted neutrophil markers (*mpx* ^1^, *lyz* ^2^, *npsn* ^3^) or macrophage/monocyte (Mϕ) markers (*mfap4* ^4^, *ccl34a.4* ^5^, *mpeg1.1* ^6^). **g)** Scaled expression of cytokines in indicated neutrophil and Mϕ clusters. **a**–**g)** Scale bars: 50 µm. Abbreviations: 4OHT, 4-hydroxytamoxifen; cht, caudal hematopoietic tissue; h, hours; hpl, hours post-lesion; hs, heat shock; Mϕ, macrophages/monocytes; sc, spinal cord; UMAP, uniform manifold approximation and projection.

**Supplementary Figure 4.**
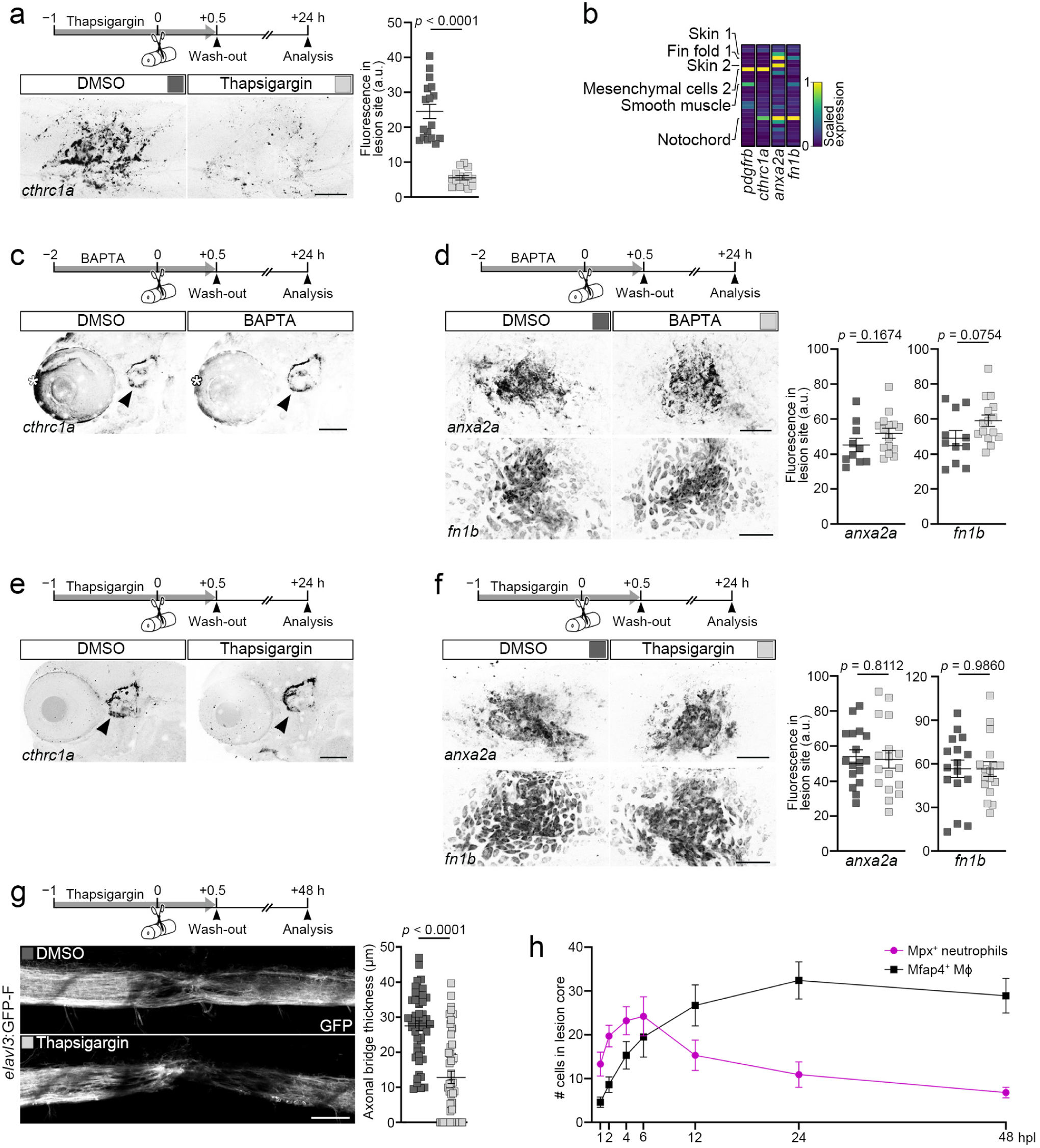
Characterization of BAPTA and thapsigargin treatments and dynamics of immune cell recruitment. **a)** Short-term treatment with the sarco/endoplasmic reticulum Ca²⁺ ATPase inhibitor thapsigargin leads to reduced *cthrc1a* mRNA expression in the lesion site at 24 hpl. Images shown are maximum intensity projections of the lesion site (lateral view; rostral is left). Data are means ± SEM. Each data point represents one animal. Two-tailed Mann–Whitney test. *n*_DMSO_ = 17, *n*_Thapsigargin_ = 14. **b)** Scaled expression of *pdgfrb*, *cthrc1a*, *anxa2a*, and *fn1b* in all major clusters. Clusters with higher expression levels are indicated. Note: *pdgfrb* and *cthrc1a* expression is highest in mesenchymal cells 2, while *anxa2a* and *fn1b* transcripts localize mainly to other cell clusters. **c)** Short-term treatment with the cell permeant Ca^2+^ chelator BAPTA-AM (abbreviated as BAPTA) does not reduce *cthrc1a* mRNA expression in the inner ear (arrowheads). Images shown are maximum intensity projections (lateral view; rostral is left). *n* = 20 for each condition. Images shown are heads of specimen presented in Fig. 4d. Asterisks indicate background signal. **d)** Short-term treatment with BAPTA does not reduce *anxa2a* and *fn1b* expression in the lesion site at 24 hpl. Images shown are maximum intensity projections of the lesion site (lateral view; rostral is left). Data are means ± SEM. Each data point represents one animal. Two-tailed Student’s *t*-test. *anxa2a*: *n*_DMSO_ = 10, *n*_BAPTA_ = 15; *fn1b*: *n*_DMSO_ = 11, *n*_BAPTA_ = 15. **e)** Short-term treatment with thapsigargin does not reduce *cthrc1a* mRNA expression in the inner ear (arrowheads). Images shown are maximum intensity projections (lateral view; rostral is left). *n*_DMSO_ = 17, *n*_Thapsigargin_ = 14. Images shown are heads of specimen presented in Supplementary Fig. 4a. **f)** Short-term treatment with thapsigargin does not reduce *anxa2a* and *fn1b* mRNA expression in the lesion site at 24 hpl. Images shown are maximum intensity projections of the lesion site (lateral view; rostral is left). Data are means ± SEM. Each data point represents one animal. Two-tailed student’s *t*-test. *anxa2a*: *n* = 17 for each condition; *fn1b*: *n*_DMSO_ = 16, *n*_Thapsigargin_ = 17. **g)** Short-term treatment with thapsigargin reduces the thickness of the axonal bridge (white) at 48 hpl. Data are means ± SEM. Each data point represents one animal. Two-tailed Mann–Whitney test. *n*_DMSO_ = 48, *n*_Thapsigargin_ = 47 over three independent experiments. **h)** Dynamics of Mpx^+^ neutrophil and Mfap4^+^ Mϕ recruitment after SCI. Neutrophil numbers peak at 4– 6 hpl in the lesion core. Mϕ numbers peak at 12 hpl and plateau until 48 hpl. Data are means ± SEM. *n* = 10 for each timepoint. **a**–**h)** Abbreviations: a.u., arbitrary unit; h, hours; hpl, hours post-lesion; Mϕ, macrophages/monocytes. Scale bars: 100 µm (**c**, **e**), 50 µm (**a**, **d**, **f**, **g**).

**Supplementary Figure 5.**
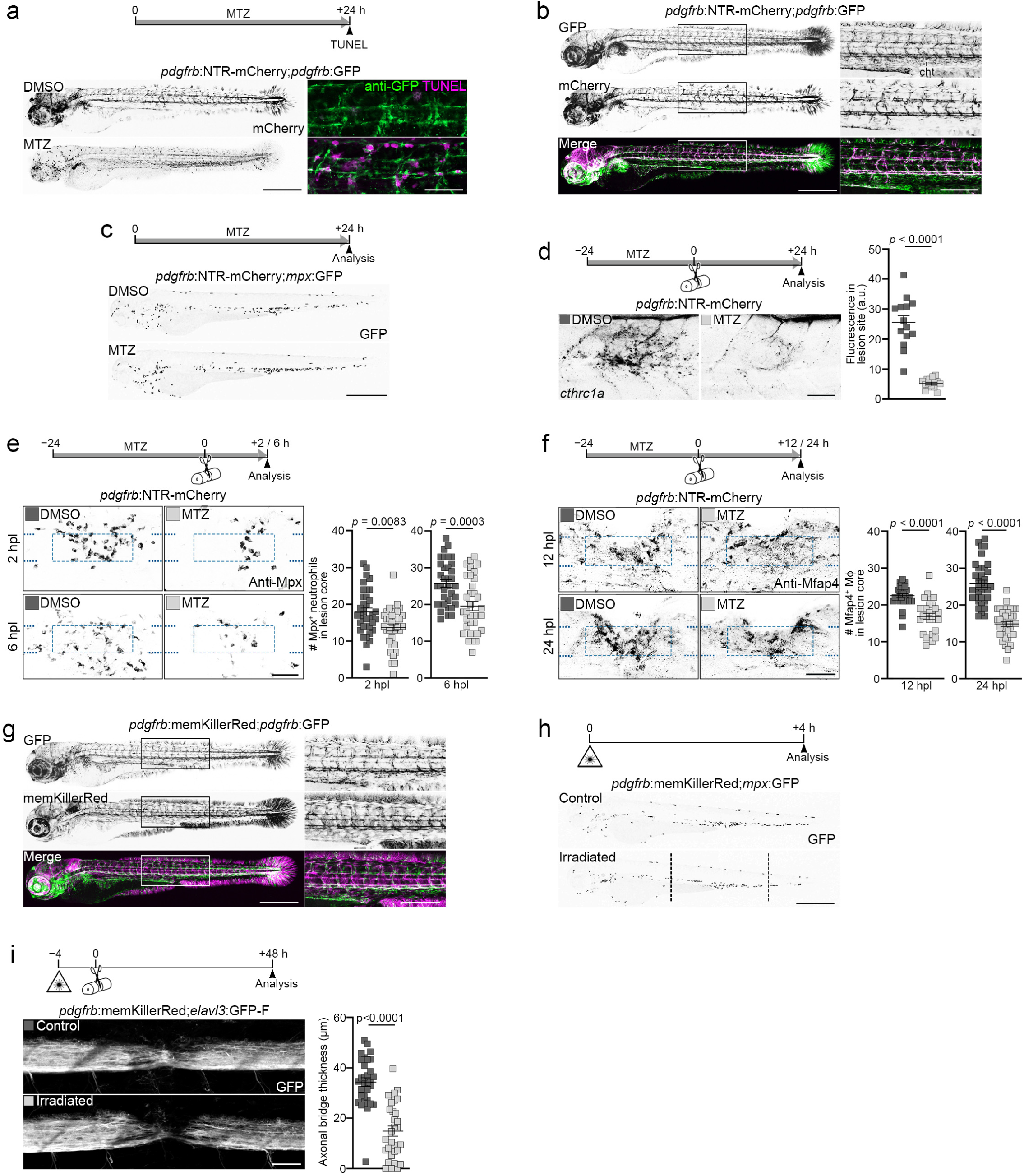
*cthrc1a*^+^ fibroblasts induce the inflammatory response. **a)** Treatment of *pdgfrb*:NTR-mCherry transgenic animals with metronidazole (MTZ) depletes transgene-expressing *pdgfrb*^+^ cells (mCherry; black) through induction of cell death (TUNEL^+^; magenta). *n* > 10 for each condition. **b)** *pdgfrb*^+^ cells co-label with mCherry and GFP in *pdgfrb*:NTR-mCherry;*pdgfrb*:GFP transgenic animals. Note: mCherry fluorescence is only detectable in a subset of GFP^+^ cells due to transgene silencing and mCherry fluorescence is largely absent from the trunk region containing the caudal hematopoietic tissue (cht). Shown is the same specimen (DMSO) as presented in Supplementary Fig. 5a. *n* > 10. **c)** Ablation of *pdgfrb*^+^ cells in *pdgfrb*:NTR-mCherry transgenic animals does not lead to the recruitment of Mpx^+^ neutrophils to the fibroblast niche. *n* = 3 for each condition. **d)** Treatment of *pdgfrb*:NTR-mCherry transgenic animals with MTZ leads to reduced *cthrc1a* mRNA expression in the lesion site at 24 hpl. Two-tailed Student’s *t*-test. *n*_DMSO_ = 14, *n*_MTZ_ = 16. **e)** Treatment of *pdgfrb*:NTR-mCherry transgenic animals with MTZ leads to reduced appearance of Mpx^+^ neutrophils in the lesion core at the indicated timepoints. 2 hpl: Two-tailed Mann–Whitney test, *n* = 35 for each condition over three independent experiments; 6 hpl: Two-tailed Student’s *t*-test, *n* = 35 for each condition over two independent experiments. **f)** Treatment of *pdgfrb*:NTR-mCherry transgenic animals with MTZ leads to reduced appearance of Mfap4^+^ Mϕ in the lesion core at the indicated timepoints. 12 hpl: Two-tailed Mann–Whitney test, *n*_DMSO_ = 30, *n*_MTZ_ = 28; 24 hpl: Two-tailed Student’s *t*-test, *n* = 35 for each condition. **g)** *pdgfrb*^+^ cells co-label with memKillerRed and GFP in *pdgfrb*:memKillerRed;*pdgfrb*:GFP transgenic animals. Shown is the same specimen as presented in Fig. 5a (Control). *n* = 15. **h)** Spatially restricted exposure of *pdgfrb*:memKillerRed transgenic animals to green light does not lead to the recruitment of Mpx^+^ neutrophils to the irradiated trunk tissue. *n* = 3 for each condition. Vertical dashed lines indicate area of irradiation. **i)** Spatially restricted exposure of *pdgfrb*:memKillerRed transgenic animals to green light reduces the thickness of the axonal bridge (white) at 48 hpl. Two-tailed Mann–Whitney test, *n*_Control_ = 31, *n*_Irradiated_ = 30 over two independent experiments. **a**–**i)** Data are means ± SEM. Each data point represents one animal. Images shown are maximum intensity projections of uninjured animal, uninjured trunk, or lesion site (lateral view; rostral is left). Dashed lines indicate location of the spinal cord, dashed square indicates the region of quantification, solid square indicates the location of the inset. Abbreviations: a.u., arbitrary unit; cht, caudal hematopoietic tissue; h, hours; hpl, hours post-lesion; Mϕ, macrophages/monocytes; MTZ, metronidazole. Scale bars: 500 µm (overview in **a**, overview in **b**, **c**, overview in **g**, **h**), 250 µm (inset in **b**, inset in **g**), 50 µm (inset in **a**, **d**–**f**, **i**).

**Supplementary Figure 6.**
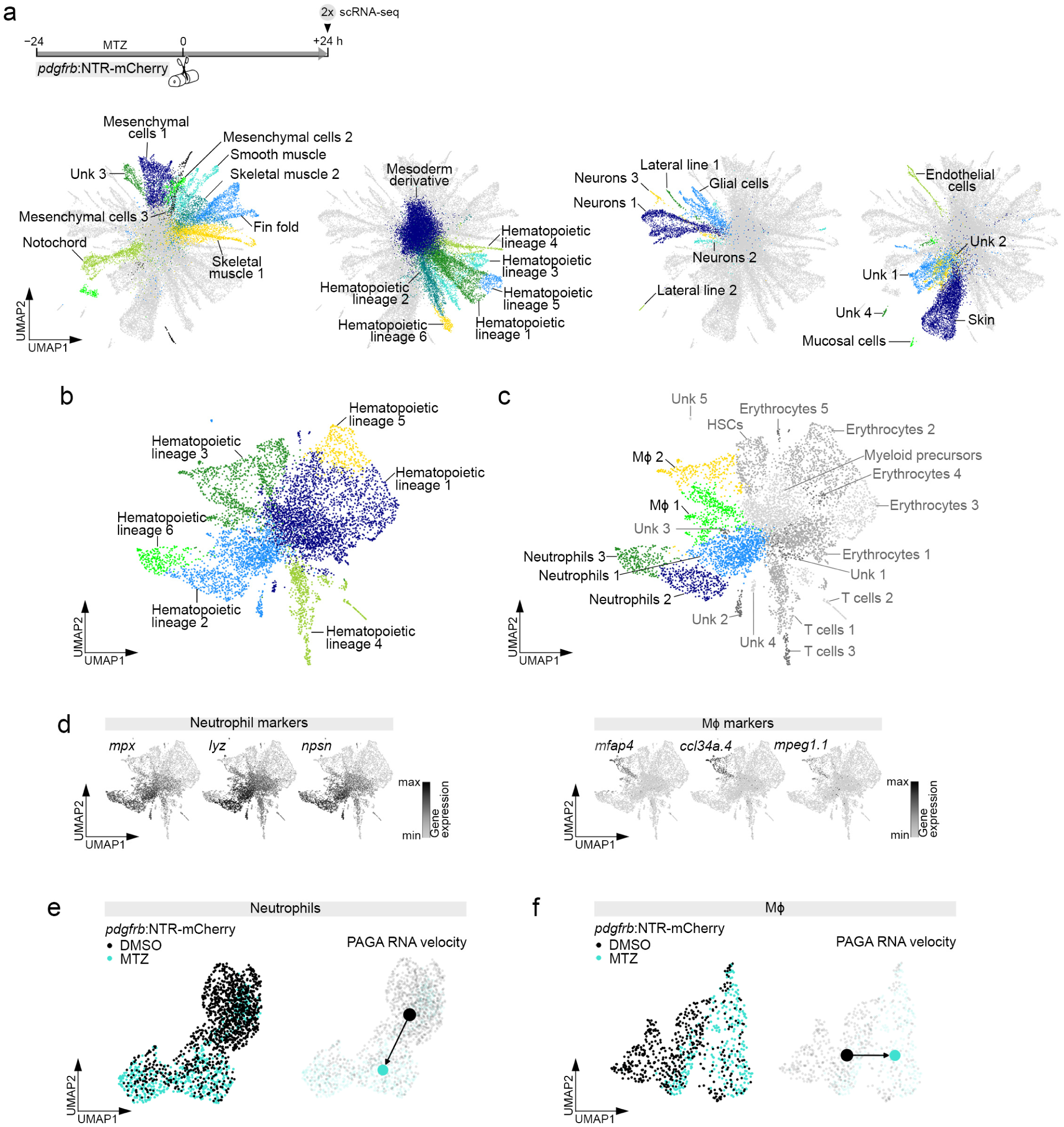
Single cell transcriptome analysis of the spinal lesion environment after *pdgfrb*^+^ fibroblast depletion. **a)** Timeline of scRNA-seq experiment and UMAP representation of the post-filtering clustering results. Cells for scRNA-seq analysis were isolated from DMSO or MTZ-treated *pdgfrb*:NTR-mCherry transgenic animals. **b)** UMAP representation of all identified hematopoietic lineage clusters. **c)** UMAP representation of all hematopoietic lineage cells after second-level clustering. Identified Mϕ and neutrophil clusters are highlighted in color. **d)** UMAP representation of all hematopoietic lineage cells showing scaled expression (black) of neutrophil markers (*mpx* ^1^, *lyz* ^2^, *npsn* ^3^) or Mϕ markers (*mfap4* ^4^, *ccl34a.4* ^5^, *mpeg1.1* ^6^). **e**–**f)** RNA velocity analysis suggests a transition of the neutrophil’s and Mϕ’s transcriptional signatures between DMSO and MTZ-treated *pdgfrb*:NTR-mCherry transgenic animals. **a**–**f)** Abbreviations: Mϕ, macrophages/monocytes; MTZ, metronidazole; PAGA, partition-based graph abstraction; UMAP, uniform manifold approximation and projection; unk, unknown.

**Supplementary Figure 7.**
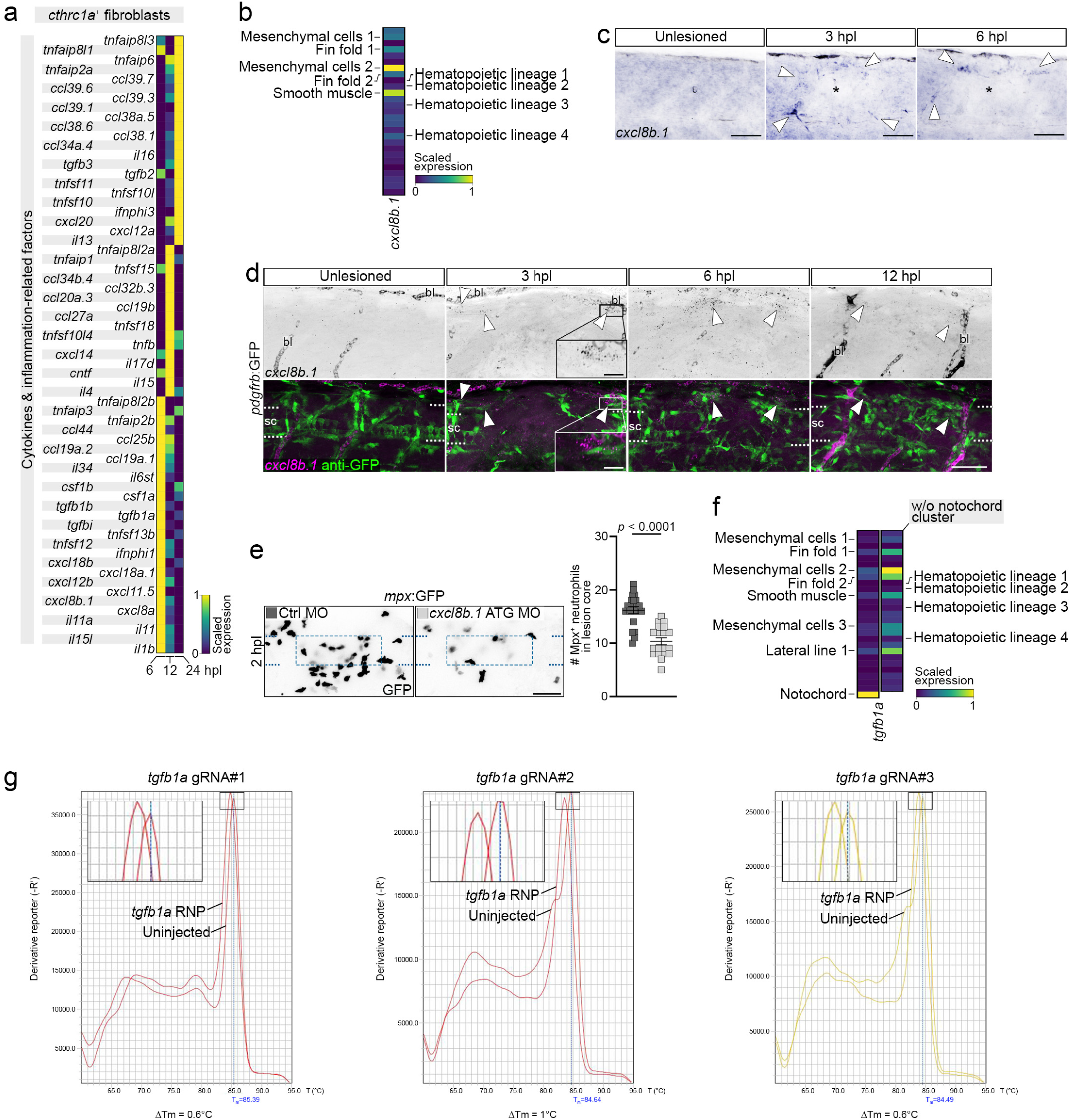
Cxcl8 and Tgfβ1 mediate the inflammation-modulating functions of *cthrc1a*^+^ fibroblasts. **a)** Scaled expression of indicated cytokines and inflammation-related factors in *cthrc1a*^+^ fibroblasts at 6, 12, and 24 hpl. **b)** Scaled expression of *cxcl8b.1* in all major clusters. Clusters with higher expression levels are indicated on the left side. Note: hematopoietic lineage clusters (indicated on the right side) show only low-level expression of *cxcl8b.1*. **c)** *cxcl8b.1* mRNA expression is upregulated at the wound margin at 3 and 6 hpl (arrowheads), as determined by *in situ* hybridization (lateral view; rostral is left). Asterisks indicate the lesion center. *n* ≥ 6 for each timepoint. **d)** *cxcl8b.1* mRNA expression (black/magenta) is upregulated in *pdgfrb*^+^ fibroblasts (anti-GFP^+^; green) at the wound margin at 3, 6, and 12 hpl. Images shown are maximum intensity projections of the lesion site (lateral view; rostral is left). Arrowheads indicate overlap of GFP^+^ cells and *cxcl8b.1* mRNA. For 6 hpl, the same specimen is shown as presented in Fig. 7d. *n* ≥ 10 for each timepoint. **e)** Delivery of a translation blocking morpholino (ATG MO) targeting *cxcl8b.1* into the zygote leads to reduced appearance of Mpx^+^ neutrophils (black) in the lesion core at 2 hpl. Data are means ± SEM. Each data point represents one animal. Two-tailed Student’s *t*-test, *n*_Ctrl MO_ = 20, *n_cxcl8b.1_* _ATG MO_ = 19. **f)** Scaled expression of *tgfb1a* in all major clusters. Clusters with higher expression levels are indicated on the left side. Note: *tgfb1a* transcript abundance is second highest in mesenchymal cells 2. Also note: hematopoietic lineage clusters (indicated on right side) show only low-level expression of *tgfb1a*. **g)** Melting curve analysis indicates efficient genome editing of selected *tgfb1a* gRNAs. **a**–**g)** Dashed lines indicate location of the spinal cord, dashed square indicates the region of quantification. Abbreviations: bl, blood; ctrl, control; h, hours; hpl, hours post-lesion; RNP, gRNA/Cas9 ribonucleoprotein complex; sc, spinal cord; Tm, melting temperature; w/o, without. Scale bars: 50 µm (**c**–**e**), 10 µm (inset in **d**).

**Supplementary Figure 8.**
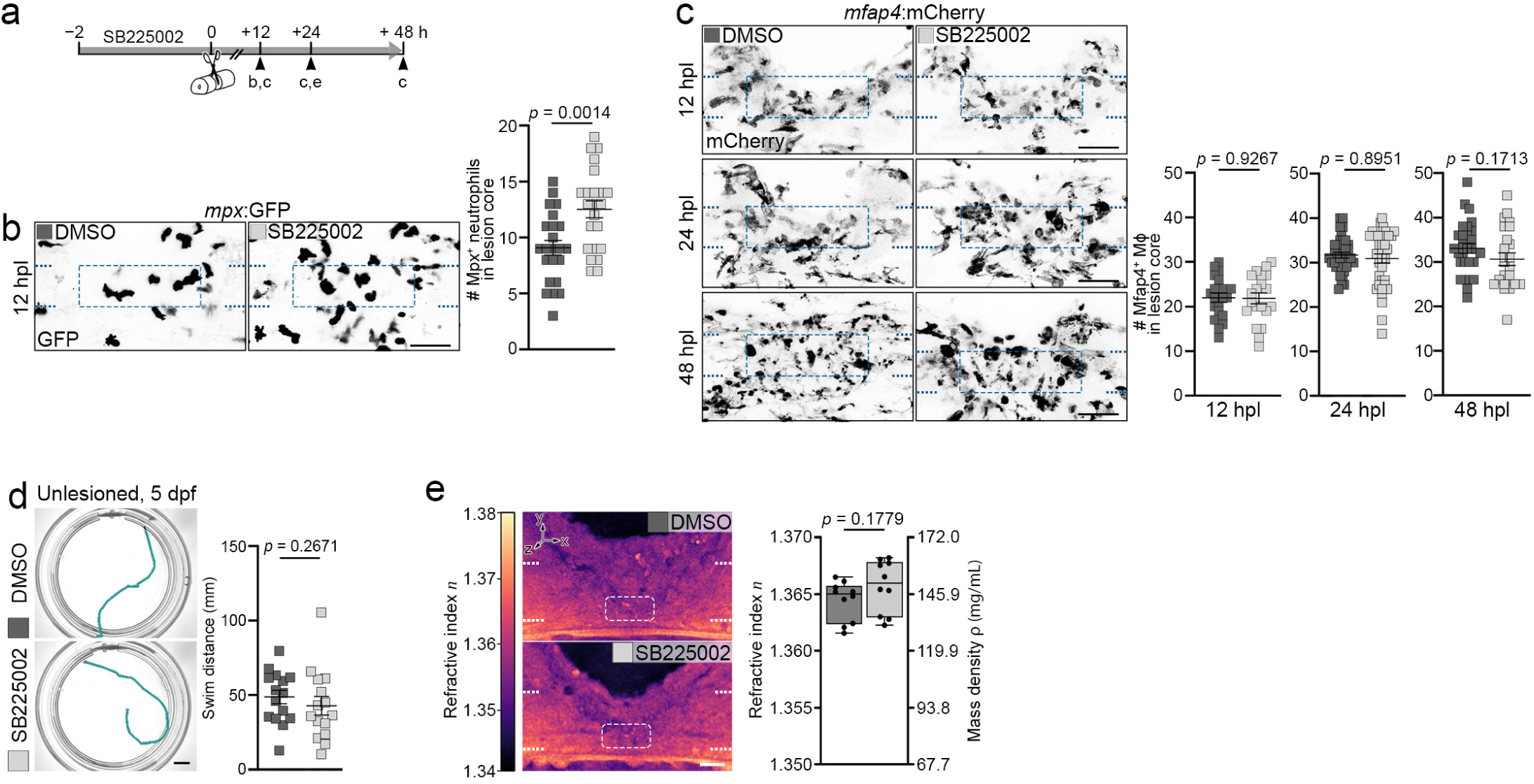
Characterizing the effect of SB225002 treatment on Mϕ and neutrophil numbers, swimming function, and refractive index distribution. **a)** Timeline for treatments with the Cxcr1/2 antagonist SB225002 shown in **b**–**e**. **b)** SB225002 treatment leads to increased retention of Mpx^+^ neutrophils (black) in the lesion core at 12 hpl. Two-tailed Student’s *t*-test, *n* = 23 for each condition. **c)** SB225002 treatment does not affect the number of Mfap4^+^ Mϕ (black) in the lesion core at indicated timepoints. 12 hpl: Two-tailed Student’s *t*-test, *n* = 20 for each condition. 24 hpl: Two-tailed Mann– Whitney test, *n* = 37 for each condition over two independent experiments. 48 hpl: Two-tailed Student’s *t*-test, *n*_DMSO_ = 28, *n*_SB2205002_ = 24. **d)** Swimming distance is not impaired in unlesioned animals after two days of SB225002 treatment. Two-tailed Mann–Whitney test, *n* = 15 for each condition. **e)** SB225002 treatment does not affect the refractive index (*n*) of the lesion site in live animals at 24 hpl, as determined using optical diffraction tomography (ODT). Two-tailed Student’s *t*-test, *n* = 10 for each condition over two independent experiments. **a**–**e)** Data are presented as mean ± SEM (**b**–**d**) or as box plots showing the median, first, and third quartiles with whiskers indicating the minimum and maximum values (**e**). Each data point represents one animal. Images shown are maximum intensity projections (**b**, **c**) or sagittal optical slices through a refractive index tomogram (**e**) of the lesion site (lateral view; rostral is left). Dashed lines indicate location of the spinal cord, dashed square indicates the region of quantification. Scale bars: 5 mm (**d**), 50 µm (**b**, **c**), 20 µm (**e**). Abbreviations: h, hours; hpl, hours post-lesion; dpf, days post-fertilization.

